# Synthetic lethality of fungal catabolite repressors reveals essential coupling of carbon repression with mitochondrial and ribosomal gene expression

**DOI:** 10.64898/2026.05.08.723686

**Authors:** Anmoldeep Randhawa, Raubins Kumar, Mayank Gupta, Syed Shams Yazdani

## Abstract

Carbon catabolite repression (CCR) enables fungi to preferentially utilize favorable carbon sources through global transcriptional control. In filamentous fungi, CCR is classically attributed to the C2H2 zinc-finger repressor Mig1/CreA; however, residual repression following Mig1 disruption suggests additional regulatory layers in *Talaromyces pinophilus* (previously called *Penicillium funiculosum* NCIM1228). Here, we identify Mig3, a previously uncharacterized, Eurotiales-specific C2H2 transcription factor that functions redundantly with Mig1 in CCR. Mig3 contains conserved N-terminal zinc-finger motifs and a fungal transcription factor middle homology region and binds canonical Mig1 response elements with higher DNA affinity. Genetic analyses reveal that simultaneous loss of Mig1 and Mig3 is synthetically lethal, indicating an essential shared function beyond carbon repression. Integrative transcriptomic, chromatin immunoprecipitation sequencing, and proteomic analyses demonstrate that Mig1 and Mig3 directly regulate genes encoding mitochondrial and ribosomal components, including key factors involved in oxidative phosphorylation, while repressing genes required for alternative carbon metabolism. Loss of either repressor compromises expression of respiratory and ribosomal genes and confers hypersensitivity to mitochondrial and ribosomal inhibitors. In addition, both repressors physically associate with mitochondrial, ribosomal, and central metabolic proteins, suggesting coordination of transcriptional and post-transcriptional regulatory mechanisms. Together, our findings redefine fungal catabolite repressors as metabolic gatekeepers that couple CCR to mitochondrial and ribosome biogenesis, thereby controlling the metabolic switch between core cellular growth processes and enzyme secretion.

## 1. Introduction

Yeasts and filamentous fungi are major biofactories for the production of many essential enzymes and metabolites in the pharmaceutical, cosmetic, food, feed, and chemical industries. These saprophytic microbes are inherently adapted to dynamic nutritional environments with mixtures of simple carbohydrates (e.g., glucose) and complex carbohydrates (e.g., lignocellulose) (1). They rely on a continuum of growth strategies to shift from the most preferred nutrient to the least preferred one based on cost-benefit trade-offs (2–4). These strategies primarily involve the regulation of gene expression in response to environmental conditions. The sequential utilization of resources is attributed to the catabolic repression of secondary metabolic pathways in the presence of a preferred source, followed by transcriptional activation of secondary metabolic pathways upon depletion of the preferred source. The change in gene expression could occur through a classical diauxic shift, a steady-state metabolic transition from the most preferred source to the secondary source, stochastic sensing, and/or bet-hedging between different metabolic states (2–5). In addition to altering the gene expression of their preexisting genetic repertoire, microbes also increase their metabolic capacity, albeit slowly, through genetic adaptation via mutations, gene deletions, duplications, etc (6–8). However, these tightly regulated intrinsic adaptations prevent microbes from achieving maximal fitness in stable growth environments such as bioreactors (8). Besides being both time-consuming and energy-demanding, these intricate regulatory systems prevent the system from operating to its maximum potential in light of impending change (2). Therefore, for a natural isolate to transform into an industrial strain, these regulatory systems are altered by genetic manipulations to maintain high-level gene expression in the stable growth conditions of a bioreactor (8).

In the budding yeast *Saccharomyces cerevisiae*, carbon catabolite repression (CCR) is largely mediated by a conserved C2H2 zinc finger transcriptional repressor called Mig1 and, to a lesser extent, by the paralogs Mig2 and Mig3 (6, 7). Similarly, *Candida albicans* has two redundant catabolite repressors, Mig1 and Mig2 (9). In the presence of glucose, Mig1 and its paralogs repress the transcription of genes encoding enzymes that utilize maltose, sucrose, galactose, and other alternative carbon sources. Mig1 and its paralogs also mediate indirect catabolite repression by repressing genes that encode transcriptional activators (e.g., Gal4) of genes encoding enzymes that utilize alternate sugars (6). Catabolite repressors of *S. cerevisiae* and *C. albicans* are functionally redundant and can be simultaneously deleted to achieve catabolically de-repressed yeast strains. Furthermore, the three paralogs could bind to the same DNA binding motif, 5′-GCGGGG-3′ or a similar DNA binding motif, 5′-SYGGRG-3′ (6). Filamentous fungi such as *Aspergillus nidulans* and *Neurospora crassa* have a C_2_H_2_ zinc finger containing the Mig1/Mig2/Mig3 ortholog called CreA/Cre-1 and bind to the conserved DNA motif, 5′-SYGGRG-3′ (10–13). Although the regulatory cascades of metabolism are well studied in model organisms such as *S. cerevisiae* and *N. crassa*, natural diversity in metabolic regulation and growth strategies in non-model industrial strains has received less attention.

Our previous bioprospecting research identified *Talaromyces pinophilus* (previously known as *Penicillium funiculosum*) NCIM1228 as a non-model fungal isolate with a high-performing lignocellulolytic secretome (14). The efficacy of its enzymatic repertoire surpasses that of the current industrial strain, *T. reesei*, suggesting that NCIM1228 could be a potential industrial strain for lignocellulolytic enzyme production (15). Genome sequencing of the isolate revealed that it harbours a transversion mutation in the Mig1 gene at the 400^th^ nucleotide (16, 17). Consequently, the *T. pinophilus* Mig1 (TpMig1) gene encodes a truncated catabolite repressor that prematurely terminates at the 134^th^ amino acid. The mutation was considered to be one of the factors contributing to the enhanced cellulolytic activity observed in the secretome of this fungal strain (15, 16). The strain was further engineered to completely disrupt the TpMig1 gene, which resulted in a 2-fold increase in the secretion and lignocellulolytic activity of its secretome (16, 18)). Furthermore, we also identified the transcriptional activator of cellulase, Clr-2 and over-expressed it to reinforce the high expression of cellulase genes (19, 20). Further, we identified the role of calcium signaling in cellulase production and overexpressed the calcium-activated kinase Ssp1 to achieve a 40% increase in cellulase production (20, 21). However, despite the disruption of the catabolite repressor TpMig1 and the overexpression of the transcriptional activator Clr-2, the engineered strain could not fully produce cellulase in the presence of glucose, unlike *T. reesei* (22). This indicates that the catabolite repression mechanisms are diverse and differentially evolved in filamentous fungi.

The present study was conducted to fully understand the dynamics of CCR in *T. pinophilus* NCIM1228. The continuance of CCR, despite the disruption of TpMig1, indicated the presence of other potent catabolite repressors in *T. pinophilus*. Bioinformatic analysis of the *T. pinophilus* genome identified another zinc finger transcription factor that shares high similarity with TpMig1 in its zinc fingers. Gene deletion analysis, followed by additional phenotypic assays, identified this transcription factor as a catabolite repressor, which we named TpMig3. Additionally, the double deletion of catabolite repressors was found to be lethal, unlike *S. cerevisiae* and *C. albicans*. Multi-omics studies of NCIM1228, Δ*mig1*, and Δ*mig3* further mapped the regulon of the repressors. Besides acting as catabolite repressors, the two catabolite repressors also function as transcriptional activators of many essential genes, including those of oxidative phosphorylation, in the presence of glucose. Their deletion significantly affects ATP synthase expression, leading to synthetic lethality associated with the loss of these two catabolite repressors.

## 2. Materials and Methods

### 2.1. Strains, growth media and culture conditions

All fungal strains used in the study are derivatives of *T. pinophilus* NCIM1228. The media used were potato dextrose (PD) broth and agar, LMP (1% malt extract, 0.05% soya peptone), SD minimal medium (2% glucose and 0.67% yeast nitrogen base without amino acids (Difco) and RCM medium (2.4% soya peptone, 0.59% KH_2_PO_4_, 0.312% (NH_4_)_2_SO_4_, 0.005% CaCl_2_·2H_2_O, 0.005% yeast extract, 2.4% Avicel, and 2.14% wheat bran). Mandel’s minimal (MM) medium (Hajny and Reese, 1969) having 4% glucose and 4% Avicel was used for RNA profiling by RNA-seq, and RT-qPCR, protein expression by western blotting, ChIP-seq and Co-IP experiments. We used PD for fungal culturing, low malt extract peptone (LMP) agar for transformations, Mandel’s medium (MM), and RCM/MM medium for cellulase production.

The growth profile on agar plates was observed by spotting 5μl of 10^4^ conidiospores/ml suspension on SC agar with a 2% carbon source followed by 48 h of incubation at 28 °C. To determine the differential growth profile in liquid medium, Mandel’s medium with 4% glucose was inoculated with primary culture at 5%, and the cultures were incubated at 28 °C for 24 h. Mycelia were collected by filtration through Mira cloth, and the collected mycelia were dried for 72 h at 70 °C and weighed

### 2.2. Gene cloning, vector construction and transformation

pBIF-*Tpmig3* deletion cassette construction: A split marker disruption cassette was designed by substituting the ORF region of *Tpmig3* with a hygromycin expression cassette. The resultant 6.1 kb homologous recombination-based *Tpmig3* deletion cassette was chemically synthesized and ligated into the T-DNA of pCambia1302 at the *Mau*BI/*Xho*I restriction sites. The resultant vector, pAR1, was transformed into NCIM1228 via *Agrobacterium* mediated method and transformants were selected in the presence of hygromycin (16). Hygromycin-resistant transformants were screened by PCR, and Δ*mig3* transformants exhibiting an amplified DNA product size corresponding to the *TpMig3* deletion cassette were selected. The Δ*mig3* transformants were also confirmed for *TpMig3* deletion by Rapid Amplification of cDNA Ends (RACE). Two hundred bases from both the 5′ and 3′ ends of the *TpMig3* ORF were amplified using RACE.

pBIF-*ble-tet_off_mig1* construction, transformation and expression: A 1527bp *Tpmig1* gene along with terminator was amplified and cloned at Nco1/HindIII site in pBIF-Nat by replacing *egfp* gene, resulting in pBIF-*nat*-*mig1*. The *gpd* promoter was replaced by a 2582 bp tet-off cassette from pFW15.1 (23) at restriction sites XmaJ1/Nco1. The hygromycin resistance cassette of *p*BIF*-nat-tet_off_-mig1* was replaced by a bleomycin resistance cassette having a homologous region upstream to Mig1 from the TpMig1^88^ deletion cassette (16) at Kpn1/XmaJ1 restriction sites to form pBIF-*ble-tet_off_mig1*. The resultant 13.6 kb pBIF-Tet-Off-mig1 *Agrobacterium*-based expression vector, having zeocin as a selection marker, was transformed to replace the native *Tpmig1* gene and promoter by homologous recombination. pBIF-Tet-Off-mig1 was transformed into NCIM1228 and Δ*mig3*, and the transformants were screened in the presence of zeocin. The stable transformants were confirmed for integration by PCR amplification of the bleomycin resistance cassette and *Tet-Off* regulated cassette. To optimize the doxycycline concentration required to switch off the expression of *Tpmig1*, the transformants were cultured in SC medium with or without varying concentrations of doxycycline for 48 hours. Transcript levels of the TpMig1 under varying concentrations of doxycycline were determined by semi-quantitative and quantitative real-time PCR. Both methods indicated that the addition of 50 µg/ml Dox was effective in downregulating the expression of TpMig1 after 48 hours

pET28a-6X*his-mig1* and pET28a-6X*his-mig3* construction: a 402bp fragment encoding TpMig1 and TpMig3 zinc fingers was amplified using Mig1 PspX1 R and Mig1 NdeI F for TpMig1, and Mig3 Xho1R and Mig3 Nco1 F for TpMig3 from the NCIM1228 genome and cDNA, respectively. The amplified fragments were cloned into pET28a(+) at *Nde*I/*Psp*X1 and NcoI/Xho1 restriction sites for Mig1 and Mig3, respectively, and the positive clones were confirmed by restriction digestion and colony PCR.

### 2.3. Homology Modelling

We next used AlphaFold and Swiss-Model to construct a homology model of the TpMig3 protein. Both AI programs predicted a similar structure of TpMig3, having a C2H2 zinc finger domain and a large fungal_TF_MHR domain. However, due to the lack of a closely related structure, both AI programs failed to predict if TpMig3 has a transmembrane domain. Next, we constructed a homology model of TpMig1 and TpMig3 using the Swiss-model AI platform. The closest template for TpMig3 was identified as a zinc finger DNA-binding motif from Zif268 (a mouse immediate early protein). The consensus DNA-binding site (PDB: 1zaa.1) was determined by X-ray crystallography at a resolution of 2.1Å and refined to a crystallographic R factor of 18.2% (Pavletich and Pabo, 1991). A template search for TpMig1 identified a consensus sequence-based Zn finger (PDB: 1mey.1) bound to DNA. To rule out any template-related variations in the Zn finger models of TpMig1 and TpMig3, models were constructed using 1zaa.1 as the template. Ramachandran statistics revealed that 96.08% of the amino acid residues of both TpMig1 and TpMig3 models were incorporated in the favoured regions of the plot (Fig. S1). While there were no outliers in the TpMig3 model, one cysteine residue at the 34^th^ position was considered an outlier in TpMig1, indicating that the models are reliable for structural analysis (Fig. S1). Both models were superimposed on the template (1zaa) to identify any peculiarities in the Zn finger domains of TpMig1 and TpMig3.

### 2.3. Glucose and cellobiose uptake assay

10^6^ spores of NCIM1228, Δ*mig1*, and Δ*mig3* were cultured in potato dextrose broth for 24 h. 10% of the primary culture was added to 50 ml of SC media containing 1% glucose or 1% cellobiose. Supernatant samples were collected at different time intervals, and residual glucose and cellobiose in the supernatant were analysed using an HPLC system (Agilent Technologies, USA) equipped with an Aminex HPX-87H anion exchange column (Bio-Rad, USA) and a refractive index detector. Areas obtained from 1 g/l standards of glucose and cellobiose (Absolute standards) were used to calculate residual glucose and cellobiose concentrations in the test samples.

### 2.4. Enzyme assays and biomass hydrolysis

Secretomes were evaluated for enzyme activities on 0.5% crystalline cellulose (Avicel, Sigma), 1% carboxymethylcellulose (CMC, Sigma), 1% xylan (Himedia), and 1% starch (Himedia) as described previously [18]. Citrate–phosphate buffer (50 mM, pH 4.0) was used for diluting the secretomes and carrying out enzyme assays. Enzymes and substrates are incubated together for 30 min (2h for crystalline cellulose) at 50 °C. The sugars released by enzyme action are measured by the dinitrosalicylic acid (DNSA) method. The absorbance at 540 nm was measured relative to a glucose or xylose standard curve. One unit of enzyme activity was defined as the amount of enzyme releasing 1 μmol of reducing sugar per min. For Biomass hydrolysis experiments, secretomes were evaluated on nitric acid-pretreated rice straw using a 20% dry biomass substrate loading concentration at enzyme concentrations of 30 g/kg dry biomass weight (DBW) (3% enzyme load). The hydrolysis reaction was conducted in 1.2-ml 96-well plates with pretreated biomass at 15% DBW loading in a 250 μl final reaction volume. The protein content of the secretomes was measured, and an appropriate volume of 30 g/kg DBW was added to the reaction mixture. The reaction was conducted in 50 mM citrate–phosphate buffer (pH 4.0) and incubated at 50 °C with constant shaking at 200 rpm for 24 h.

### 2.5. Expression of 6X-His tagged-Mig1 and Mig3 in *E. coli* and protein purification

pET28a-*6Xhis-mig1* and pET28a-*6Xhis -mig3* were transformed into BL21, and expression of 6X*-mig1* and 6X*-mig3* (∼18kD) was induced by adding 0.5 mM IPTG to the log phase cultures, and cells were harvested after 3 hours. Cells were lysed, and lysates from both uninduced and induced cultures were electrophoresed on a 12% SDS-PAGE gel (Fig. S3b, e). Western blotting with commercial rabbit anti-His antibody (Sigma) and anti-rabbit secondary HRP-conjugated secondary antibody confirmed the expression of 6X-Mig1and 6X-Mig3 in the electrophoresed samples (Fig. S3c, f). While we could achieve purified 6XHis-Mig3 by Ni^2+^-NTA affinity chromatography, 6XHis-Mig1 with >90% purity was obtained by combining Ni^2+^-NTA affinity chromatography, Cation exchange chromatography, and gel permeation chromatography (Fig. S3g, h). Ni^2+^-NTA affinity-purified Mig1 and Mig3 were dialyzed using an 8kDa cut off membrane, 50mM Sodium Phosphate buffer (SPB), pH 6.8. Dialyzed Mig3 was of >90% purity and was used for antibody generation in Balb/c mice. Dialyzed Mig1 was subjected to SP-sepharose (strong cation exchanger) purification. 1ml of SP-Sepharose slurry was washed and equilibrated with 50mM SPB, and Mig1 protein was passed through the column 5 times, and the column was washed with 50mM SPB before eluting in increasing concentration of NaCl (0.1-1M) (Fig. S3g). The cleaner elutes of Mig1 were pooled and subjected to gel permeation chromatography. Superose 200 pg 16/200 column having a column volume of 120 ml, was used. The column was equilibrated with 50mM SPB having 0.15M NaCl. Approx. 10 mg/ml of protein in 0.5 ml was injected on the column, and 128 fractions of 0.5ml size were collected. Fractions having purified Mig1 were pooled, and purified-his-tagged Mig1 was used for antibody generation in Balb/c mice (Fig. S3h).

### 2.6. Antibody generation in Balb/c mice

600 µl (∼50 mg) of purified 6XHis-Mig1 and 6XHis-Mig3 were mixed with 600 µl of Freund′s Adjuvant, complete (Sigma) and injected intra-peritoneally into Balb/C mice. The mice were further immunized with booster doses on the 21^st^ and 42^nd^ days. The booster doses were prepared by mixing 600 µl of purified 6XHis-Mig1 and 6XHis-Mig3 (∼50mg) with 600 µl of Freund′s Adjuvant, incomplete (Sigma). The blood was extracted on the 52^nd^ day of the immunization, and serum was collected. ELISA was performed to check the titer of the antibodies in the mice’s sera. A flat-bottom microtiter plate was coated with recombinant 6X-Mig1 and 6X-Mig3 protein at 4°C overnight. The plate was washed with phosphate buffer saline (PBS) three times to remove free protein. 200 µl of 1% skimmed milk in PBS was used to block any free sites in the wells for 2 h at 37°C. Test sera having antibodies were serially diluted in 0.25% skimmed milk in PBS and incubated with immobilized proteins overnight at 4°C. Pre-immune serum was diluted to 1:200 and used as a control. After three washes with PBS, HRP-conjugated goat anti-mouse IgG antibodies (Sigma) diluted to 1:10,000 were added to the wells and incubated for 1.5h at 37°C. ELISA was developed at RT by adding 100µl of 0.1mg/ml o-phenyl diamine dihydrochloride (Sigma) as chromogen and H_2_O_2_ as substrate. The reaction was stopped with 2N H_2_SO_4_. Absorbance was measured at 490nm by Spectramax M2. ELISA end point titers were 1.5times higher than the background at 10,000 dilutions of pooled Anti-Mig1 and anti-Mig3 sera. Anti-Mig1 and Anti-Mig3 were purified from sera by protein A/G affinity column chromatography (Fig. S3i)

### 2.7. Western blotting

Secretome was obtained by centrifuging the culture at 8000 rpm for 10 min, followed by buffer exchange using citrate–phosphate buffer (pH 4.0) with the help of a 3 kDa cut-off membrane. The sample was used to estimate the protein by BCA (Bicinchoninic acid) as well as to visualize protein bands on 10% SDS-PAGE (sodium dodecyl sulphate–polyacrylamide gel electrophoresis) gel. The mycelia harvested from the fungal culture were used to extract protein by the bead lysis method (20), and protein was estimated using the BCA method with bovine serum albumin as a standard. TpMig1 and TpMig3 expression in the NCIM1228 and deletion strains was detected by western blotting using mouse anti-Mig1 and mouse anti-Mig3 polyclonal antibodies and rabbit anti-mouse HRP-conjugated secondary antibody (Cell Signaling Technology).

### 2.8. Transcriptomic studies by RNA-seq

NCIM1228, Δ*mig1*, and Δ*mig3* were cultured in MM medium-4% glucose and MM medium-4% Avicel. Total RNA was isolated from log phase cultures by a Qiagen Plant Mini RNeasy kit according to the manufacturer’s instructions. Traces of DNA, if any, are removed by DNase treatment before proceeding with RNA sequencing. RNA-seq was carried out on a HiSeq 2000 platform with 125 × 2 paired-end read chemistry (Bionivid Technology Pvt Ltd). Biological replicate sequencing libraries for both strains were created with poly-A-tailed mRNA enrichment using the standard Illumina TruSeq mRNA RNA-seq protocol. The RNA-Seq reads obtained were assembled using Trinity with a reference genome-guided approach. Assembled transcripts are quantified by mapping generated sequencing reads to the assembled transcripts using the alignment mapping program Bowtie2, and alignments are coordinate-sorted by SAMtools. The quantitative program RSEM generates fragments per kilobase of transcript per million mapped reads (FPKM) from the quantitated data. The protein domains were predicted using InterProScan. For differential expression profiling, all FPKM values were normalized to the library size using the R package Edge R. The obtained p-values were plotted against log_2_ fold change using Volcano R, and volcano plots were obtained to assess the significance of up- and downregulation of transcripts, as shown in the respective figures. The DNA-binding and calcium-activated proteins were filtered and identified from the InterPro results. Heatmaps were generated from log_2_ FPKM values of selected transcription factors and calcium-activated proteins by performing hierarchical clustering using the Euclidean distance matrix option of the Morpheus heatmap tool.

### 2.9. Expression profiling by real-time qPCR

For real-time PCR experiments, mycelia were harvested from log phase cultures, and total RNA was extracted using an RNeasy kit (Qiagen). Traces of DNA were removed by DNase (Invitrogen) treatment prior to cDNA synthesis, and the RNA concentration was measured by a NanoDrop. Equal amounts of RNA were used to synthesize cDNA with an Invitrogen cDNA synthesis kit. cDNA was used as a template to carry out qRT-PCR using iTaq™ Universal SYBR® Green Supermix (Bio-Rad) and a Bio-Rad CFX96 qPCR detection system. qRT-PCR was performed in biological triplicates with tubulin as the endogenous control. Relative expression levels are normalized to tubulin, and fold changes in RNA level are the ratios of the relative expression level of the test strain to the control strain under repressing conditions and cellulase-inducing conditions to carbon repressing conditions.

### 2.10. Electrophoretic Mobility Shift Assay

For EMSA, biotinylated-labelled primers were used to amplify the DNA probe. 50 ng of recombinant zinc finger domains of TpMig1 and TpMig3 were incubated with 2.5 µM biotinylated 20-bp DNA probe in the presence/absence of varying concentrations of unlabelled probe (500 to 1000 µM), followed by 10% PAGE. Electrophoresed Biotin end-labelled duplex DNA was transferred to a nylon membrane, UV cross-linked, probed with streptavidin-HRP conjugate and incubated with the substrate to detect any retardation due to DNA-protein binding. We used 100 ng of BSA as a control, and the Biotinylated 20-bp DNA probe incubated with BSA didn’t exhibit retardation. However, 50 ng of recombinant TpMig1 and TpMig3 zinc fingers retarded the mobility of the labelled DNA motif, and increasing protein concentration from 50 ng to 100 ng in the reaction mixture further retarded the probe mobility. Furthermore, when 500 to 1000 µM of unlabelled DNA was added to the reaction mixture, it competed with the labelled probe and enhanced probe mobility. The gels were electrophoresed, and DNA was detected using the Thermofisher Light Shift Chemiluminescent EMSA Kit. We also used three additional DNA probes from the *cbh1* promoter of length 120 bp and ∼500bp corresponding to region -480 to -600 bp (P1), -1 to - 488 bp (P2) and -488 to -960 bp (P3) region upstream to gene ORF.

### 2.11. Biolayer interferometry (BLI)

To determine DNA binding affinities of the two zinc finger domains, we carried out a basic binding/dissociation assay using biolayer interferometry. BioForte OctetRED96e system was used for conducting kinetics of recombinant TpMig1 and TpMig3 Zn finger domain binding to biotinylated DNA duplex probe having sequences 5’-GGAACAAACCCCGCATATGG-3’ and 5’-CCATATGCGGGGTTTGTTCC-3’. We immobilized the biotinylated DNA probe onto the streptavidin columns and used varying dilutions of the proteins to determine a single pair of rate constants for each protein exhibiting net binding (ka) and dissociation (kd). The rate constants were used to calculate equilibrium dissociation constants (K_D_) for both TpMig1 and TpMig3.

### 2.12. Genetic crosses and Non-Random spore analysis

Δ*mig1* and Δ*mig3* were intercrossed by a modified crossing procedure (contributed by A. J. F. Griffiths, FGSC) in test tubes having SC-cross medium (KNO_3_ 4 g/l, KH_2_PO_4_ 4 g/l, MgSO_4_ .7H_2_O 4.1g/l, CaCl_2_ 0.25g/l, NaCl 0.4 g/l, trace element solution 0.4 ml, and Biotin 10 mg/ml) and a 6 cm x 4 cm spill of Whatman filter paper folded lengthwise into a “W”. Conidia from both parents were added evenly over the paper. The crosses should be incubated at 25 °C for 20 days. For non-random spore analysis, recovered ascospores were suspended in water at a spore count of 10^6^/ml and activated by heat shock at 80 °C for 2 hours to kill conidia. Equal amounts of 2X-PD broth were added to the ascospore suspension, and spores were incubated at 25 °C for twelve hours to allow them to germinate. The spore suspension was then plated onto the selection medium containing hygromycin, zeocin or both and incubated at 28 °C until mycelial growth appeared.

### 2.13. ChIP sequencing

For chromatin immunoprecipitation (ChIP), Primary cultures of *Talaromyces pinophilus* NCIM1228 were grown in potato dextrose (PD) broth for 24 h at 30°C with shaking. Mycelia were harvested by filtration, washed with sterile water, and used to inoculate secondary cultures in yeast nitrogen base (YNB) medium supplemented with 2% (wt/vol) glucose. Cultures were grown for 24 h, after which mycelia were collected by filtration through Miracloth, washed in phosphate-buffered saline (PBS), and cross-linked in PBS containing 1% formaldehyde for 30 min at room temperature with gentle agitation. Cross-linking was quenched by the addition of glycine to a final concentration of 10 mM and incubation for 20 min. Cross-linked mycelia were collected by filtration, washed three times with cold PBS, lyophilized, and stored at −80°C until further processing. Frozen mycelia were pulverized under liquid nitrogen and resuspended in a lysis buffer (Randhawa et al., 2024b). Cell lysis was performed using bashing bead tubes (Zymo Research) with alternating cycles of bead beating and incubation on ice (25 cycles of 1 min each). Lysates were sonicated to shear chromatin to an average fragment size of ∼200–500 bp, as verified by agarose gel electrophoresis. Chromatin was immunoprecipitated overnight at 4°C using in-house anti-TpMig1 or anti-TpMig3 antibodies, and immune complexes were captured using Protein A/G magnetic beads following established protocols (Ferraro and Lewis, 2018). Following reversal of cross-links and DNA purification, recovered DNA was subjected to end repair, adaptor ligation, and library amplification using the NEBNext Ultra II FS DNA Library Prep Kit (New England Biolabs).

Sequencing and primary data processing were performed by Profacgen. Libraries were sequenced on an Illumina HiSeq platform to generate 50 bp single-end reads or 150 bp paired-end reads, yielding 12-19 million reads per library. Adapter trimming and quality filtering were performed using fastp (v0.19.11), and read quality was assessed using FastQC (v0.11.5). Filtered reads were aligned to the in-house *T. pinophilus* reference genome using BWA mem (v0.7.12) with default parameters, resulting in >99% uniquely mapped reads and a duplication rate of 25% (anti-Mig1) and 30% (anti-Mig3). Fragment size distributions estimated by MACS2 indicated an average fragment length of 148 bp. The fraction of reads in peaks (FRiP) was 4.14 % for TpMig1 and 2.16 % for TpMig3 libraries.

Peaks were called using MACS2 (v2.1.0) with matched input controls, using parameters – qvalue 0.05 –call-summits –nomodel –extsize [150], resulting in 331 and 392 TpMig1 binding sites across the two biological replicates, and 320 and 129 TpMig3 binding sites across replicates. For TpMig1, 159 peaks were shared between replicates, corresponding to 48.0% reciprocal overlap and a Jaccard index of 0.353 (159 shared peaks across 450 merged intervals). For TpMig3, 59 peaks were shared between replicates, corresponding to 45.7% reciprocal overlap and a Jaccard index of 0.192 (59 shared peaks across 307 merged intervals). These levels of concordance are consistent with reproducible transcription factor ChIP-seq datasets in filamentous fungi and support the robustness of peak identification.Peak annotation relative to genomic features and transcription start sites (TSS) was performed using ChIPseeker and PeakAnnotator. The proportion of peaks located within ±1 kb of TSS was 61.7% for TpMig1 and 67.4% for TpMig3. De novo motif discovery was conducted using HOMER, MEME, and DREME, and enriched motifs were annotated using Tomtom. Functional enrichment of ChIP-seq target genes was performed using GOseq and topGO for Gene Ontology analysis and KOBAS for KEGG pathway enrichment.

### 2.14. Deletion Sensitivity Profiling

10^6^ spores of NCIM1228, Δ*mig1*, and Δ*mig3* were cultured in PD broth for 24 hours. MM medium-4% glucose, having antifungals at EC_50_ (21), was inoculated with 5% inoculum and incubated at 30 °C, 120 rpm for 24 hours. The mycelial weight was collected by filtering with Mira cloth and dried at 50 °C until a constant dry mycelial weight was obtained.

### 2.15. Co-IP LC-MS/MS

NCIM1228, Δ*mig1* and Δ*mig3* cultured in YNB Glucose for 24 hours were lyophilised, and whole mycelial protein was isolated and stored at -80 °C. For IP-Mig1 and Co-IP, mycelial protein extracts of NCIM1228 and Δ*mig1* were incubated with Anti-Mig1 overnight at 4 °C. The ProteinA/G conjugated magnetic beads were added to the samples and incubated at RT for 3h on a rocker shaker. The beads were collected and washed thrice with lysis buffer, and bound proteins were eluted in 0.1% (v/v) formic acid before proceeding for absolute quantification of all proteins by LC-MS/MS analysis by Thermofisher Orbitrap Fusion™ (21). The same procedure was also followed for For IP-Mig3 and Co-IP, and IP-preimmune sera (control).

## 3. Results

### 3.1. Partial alleviation of catabolite repression upon *Tpmig1* deletion suggests additional regulatory components

In our previous study, we identified TpMig1, a partially functional homolog of the canonical carbon catabolite repressor Mig1/CreA, in *Talaromyces pinophilus* (16). Deletion of the *mig1* gene enhanced secretory capacity under de-repressing conditions, with the Δ*mig1* strain exhibiting approximately twofold higher lignocellulolytic activity than the parental strain when grown on cellulose. However, despite this increased secretion potential, cellulase production remained constrained in the presence of glucose, indicating incomplete carbon derepression. To assess whether forced activation of the cellulolytic regulatory program could bypass this residual repression, we constitutively overexpressed the cellulase transcriptional activator Clr2 under the strong *gpd* promoter in the Δ*mig1* background. Δ*mig1* and *gpd-clr2/*Δ*mig1* strains were cultivated in media containing 4% glucose, 2% glucose plus 2% cellulose, or 4% cellulose, and lignocellulolytic activity in the secretome was quantified after five days using a filter paper unit assay. Neither strain produced detectable cellulolytic activity in glucose-rich medium, and enzyme activity in mixed glucose–cellulose cultures reached less than 50% of that observed under fully derepressing cellulose conditions (Fig. 1a). Unlike *T. reesei* (Mary Mandels and Elwyn T. Reese, 1957; Lo *et al.*, 2010; Li *et al.*, 2017), constitutive expression of the transcriptional activator *Clr2* failed to induce cellulase production in the derepressed Δ*mig1* strain of *T. pinophilus*. Additionally, cellulose could not stimulate maximum cellulase production in the presence of glucose. These findings suggest that *TpMig1* deletion only partially alleviated carbon repression in *T. pinophilus*, indicating that its carbon repression mechanisms differ from those of *T. reesei*.

**Fig. 1.**
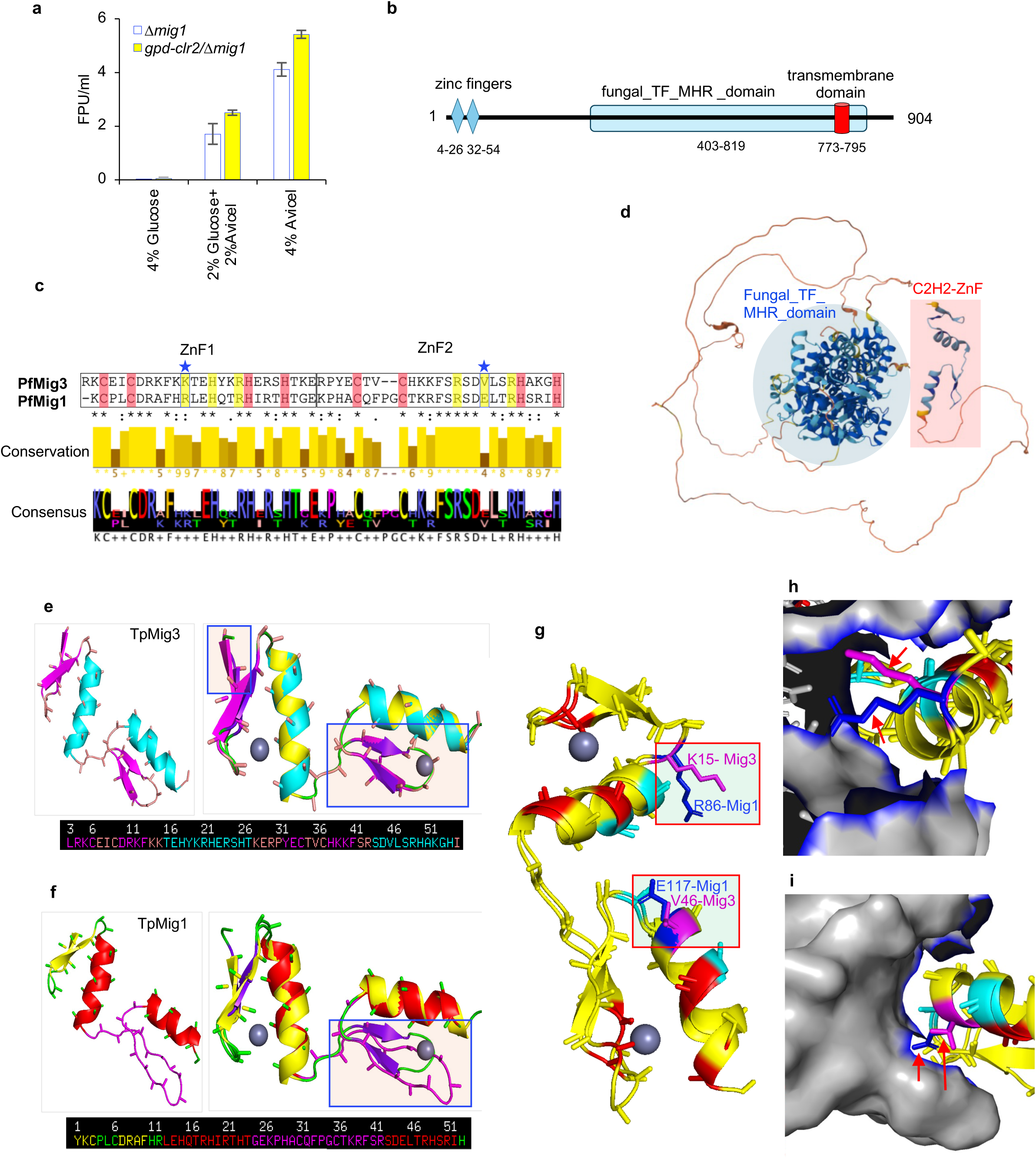
Identification of a putative catabolite repressor, TpMig3. **a,** Total cellulase activity measured by Filter Paper Unit (FPU) Assay of secretome collected from cultures of Δ*mig1* and *_PGPD_Clr-2/*Δ*mig1* incubated in different media. **b,** schematic representation of domains found in putative catabolite repressor, TpMig3. **c,** sequence alignment of zinc finger domains of TpMig1 and TpMig3, amino acids highlighted in pink are those involved in binding to zinc while those highlighted in yellow are involved in DNA binding. **d,** 3D homology model of TpMig3 prepared by SWISS-MODEL. **e,** superimposed model of zinc finger of ScMig1 and TpMig3; the boxes highlight the b-sheets and any deviation from ScMig1. **f,** superimposed model of zinc finger of ScMig1 and TpMig1; the boxes highlight the b-sheets and any deviation from ScMig1. **g,** superimposed model of zinc finger of TpMig1 and TpMig3; the boxes highlight the DNA binding amino acids in the two proteins. **h,** superimposed model of zinc finger of TpMig1 and TpMig3 with DNA showing K15 (TpMig3) and R86 (TpMig1) interact differentially at major groove of DNA. **i,** superimposed model of zinc finger of TpMig1 and TpMig3 with DNA showing V46 (TpMig3) and E117 (TpMig1) interact differentially at minor groove of DNA.

To identify additional CCR regulators, we searched the *T. pinophilus* genome using *S. cerevisiae Mig1* protein as a query. In addition to TpMig1, six genes encoding C2H2 zinc-finger transcription factors were identified (Table 1). Four corresponded to previously characterized regulators involved in conidiophore and ascospore development, stress response, or amino acid metabolism, whereas one uncharacterized gene (gene ID 29_0.82) exhibited the highest similarity to the zinc-finger domain of ScMig1 aside from TpMig1 itself. Gene 29_0.82 encodes a 904-amino-acid protein containing two N-terminal C2H2 zinc-finger motifs and a large central fungal transcription factor middle homology region (fungal_TF_MHR) (Fig. 1b). No sequence similarity to TpMig1 was detected outside the zinc-finger domain. Alignment of the zinc-finger regions revealed conservation of the Cys and His residues required for Zn coordination, but divergence at residues implicated in DNA base recognition, including substitutions within the canonical RHR and RER motifs (Fig. 1c). Based on its predicted domain architecture and similarity to Mig1 zinc fingers, we designated this protein TpMig3.

**Table 1.** Putative ZnF transcription factors showing homology to zinc finger domains of ScMig1. Using ScMig1 as a query, a Blast search of the in-house genome database was conducted to identify additional catabolite repressors in *T. pinophilus*. The transcription factor highlighted in red is named TpMig3 and represents the next best match, showing similarity to Sc Mig1 after TpMig1.

| Protein ID/<br>NCIM1228<br>Gene ID | Query<br>coverage | %<br>Identity | Name | Length<br>(AA) | Zinc<br>fingers | Other domains |
| --- | --- | --- | --- | --- | --- | --- |
| KAF3396929<br>/ 60_0.0 | 17% | 59.55% | CreA/TpMig1 | 415 | 77-97;<br>105-127 |  |
| KAF3399241<br>/ 29_0.82 | 14% | 51.92% | Transcriptional<br>regulator | 904 | 5-26;<br>34-54 | Fungal_TF_MHR:<br>403-819 |
| KAF3399308<br>/ 19_3.93 | 13% | 41.07% | hypothetical protein<br>DPV78_006976 | 345 | 280-302;<br>310-332 |  |
| KAI7977576/<br>16_0.2 | 13% | 45.59% | Master regulator of<br>conidiophore<br>development BrLA | 415 | 306-328;<br>336-359 |  |
| KAF3398996<br>/ 6_4.119 | 12% | 42.86% | C2H2 finger domain<br>transcription factor<br>sebA | 602 | 477-498;<br>504-526 |  |
| KAF3407613<br>/ 44_0.6 | 10% | 58.18% | Transcription factor<br>steA | 703 | 569-591;<br>599-619 | STE like TF domain<br>70-178 |
| KAF3400996<br>/ 53_0.81 | 10% | 37.5% | Regulator of arginine<br>metabolism | 1177 | 61-81;<br>89-110 | MADS Box TF<br>domain 257-510<br>fungal_trans 515-793 |

Homology model for the full-length TpMig3 could provide little information other than the placement of zinc finger domains and fungal_MHR_domain (Fig. 1d). Homology modelling of the zinc-finger domains of TpMig1 and TpMig3 revealed conserved overall C2H2 architecture but notable differences in local secondary structure and predicted DNA-contact residues. Superposition of the modeled zinc-finger domains with that of template (ScMig1 zinc finger) showed that while the spatial arrangement of Zn-coordinating residues was preserved, TpMig3 Zn finger has a longer β-sheet before the first Zn finger (Fig. 1e), and the Zn finger of TpMig1 exhibited a loop in place of the β-sheet before the second Zn finger (Fig. 1f). Furthermore, superposition of TpMig1 and TpMig3 Zn fingers displayed minor structural changes due to substitutions at key positions involved in base-specific interactions (Fig. 1g). In silico analysis suggested that replacement of arginine (R86) with lysine (K15) and glutamate (E117) with valine (V46) residues in the RHR and RER motifs of TpMig3 may reduce electrostatic repulsion and stabilize protein–DNA interactions relative to TpMig1 (Fig. 1h, i). Although these models do not resolve binding specificity directly, they support the idea that TpMig3 and TpMig1 recognize related DNA elements with distinct binding properties.

To examine the evolutionary distribution of Mig3, we performed BLAST searches across the fungal kingdom using TpMig3 as a query. Mig3 orthologs were identified exclusively within the order Eurotiales, predominantly in species of *Talaromyces*, *Aspergillus*, and *Penicillium*, with limited representation in a few thermophilic and saprophytic genera. Phylogenetic analysis of Mig3 homologs from 50 representative species revealed multiple lineage-specific clades, with TpMig3 grouping among recently diverged *Talaromyces* species (Fig. S2). These findings indicate that Mig3 is a relatively recent evolutionary addition to the CCR network in filamentous fungi.

### 3.2 TpMig3 functions as a carbon catabolite repressor in *T. pinophilus*

To determine the function of TpMig3, we generated a Δ*mig3* deletion strain by replacing the *mig3* open reading frame with a hygromycin resistance cassette. Correct deletion was confirmed by PCR and RACE analysis (Fig. 2a–c). Deletion of *mig3* resulted in growth phenotypes closely resembling those of Δmig1. On solid media containing glucose, potato dextrose, xylan, or Avicel, Δ*mig3* exhibited growth patterns indistinguishable from Δmig1 and distinct from the parental strain NCIM1228 (Fig. 2d). In liquid synthetic complete medium containing mixed carbon sources (2% glucose and 2% cellobiose), both Δ*mig1* and Δ*mig3* accumulated approximately 50% more mycelial biomass than NCIM1228 within 24 h (Fig. 2e). Consistent with this increased growth rate, glucose uptake in Δ*mig3* was elevated approximately 1.75-fold relative to the parent strain, matching the phenotype observed for Δ*mig1* (Fig. 2f). To directly assess carbon catabolite repression (CCR), we employed allyl alcohol (AA) and 2-deoxyglucose (2-DG) assays. In glucose-rich medium, Δ*mig3* displayed marked sensitivity to AA, indicative of de-repressed alcohol dehydrogenase expression and impaired CCR, similar to Δ*mig1* (Fig. 2g). Conversely, while growth of NCIM1228 was strongly inhibited by 2-DG in the presence of alternative carbon sources, both Δ*mig1* and Δ*mig3* grew normally (Fig. 2h), confirming defective CCR in the absence of either repressor. We next examined whether deletion of *mig3* relieved CCR on cellulase production. When cultivated in Mandel’s medium containing 2% glucose and 2% Avicel, Δ*mig3* secretomes exhibited increased total protein levels compared to NCIM1228, although to a lesser extent than Δ*mig1* (Fig. 2i). Measurement of total cellulase activity using the filter paper unit (FPU) assay revealed intermediate activity in Δ*mig3* relative to NCIM1228 and Δ*mig1* (NCIM1228 < Δ*mig3* < Δ*mig*1; Fig. 2j). Consistent with this, transcript levels of major cellulase genes were elevated more than fivefold in Δ*mig3* relative to NCIM1228 after 48 h of growth in glucose-containing medium (Fig. 2k). Under cellulase-inducing conditions (RCM medium), Δ*mig3* again displayed enhanced secretory capacity relative to the parent strain, with approximately 1.5-fold higher total secreted protein, but lower levels than Δ*mig1* (Fig. 2l; Table 2). A similar intermediate phenotype was observed for total cellulase activity and saccharification efficiency on acid-pretreated rice straw biomass (Fig. 2m, n). Together, these results demonstrate that TpMig3 functions as a bona fide catabolite repressor, whose deletion phenocopies Δ*mig1* across multiple CCR-associated traits. The intermediate derepression observed in Δ*mig3* relative to Δmig1 suggests overlapping but non-identical regulatory contributions of the two repressors to carbon metabolism and enzyme secretion.

**Fig. 2.**
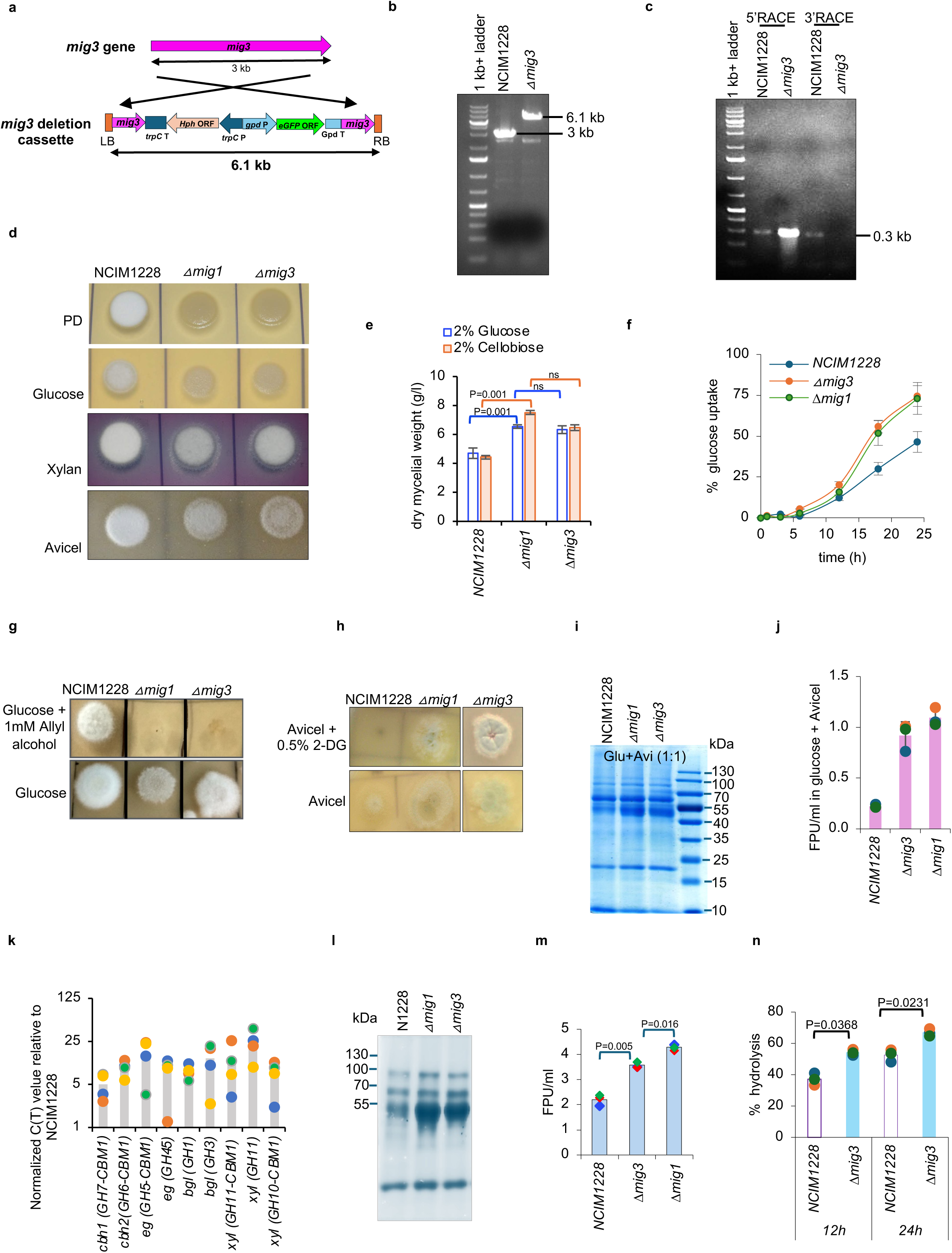
TpMig3 is a catabolite repressor. **a,** schematic showing the replacement of the TpMig3 gene with the *TpMig3* deletion cassette. **b,** PCR showing deletion of *TpMig3* in transformant, where the original gene in NCIM1228 is replaced by the Mig3 disruption cassette. **c,** RACE experiments showing that the amplification of the region at the 5’ and 3’ end to *mig3* mRNA in NCM1228 and Δ*mig3*. **d,** growth profile of NCIM1228, Δ*mig1*, and Δ*mig3* on different carbon sources. **e,** the dry mycelial weight of NCIM1228, Δ*mig1*, and Δ*mig3* when 10^6^ spores were grown in 50 ml PD broth Δ*mig3* measured by HPLC. **f,** glucose uptake by NCIM1228, Δ*mig1*, and Δ*mig3*. **g,** growth profile of NCIM1228, Δ*mig1*, and Δ*mig3* on allyl alcohol, and **h,** on deoxy-glucose. **i,** SDS-PAGE profile and (**j**) total cellulase activity of the secretome of NCIM1228, Δ*mig1*, and Δ*mig3* cultured in 2% glucose+ 2% Avicel. **k,** transcript levels of cellulase quantified in real-time by qPCR in the secretome of Δ*mig3* relative to NCIM1228. **i,** SDS-PAGE profile and (**j**) total cellulase activity of the secretome of NCIM1228, Δ*mig1*, and Δ*mig3* cultured in cellulase production media. **k,** percentage hydrolysis of acid-pretreated rice straw by NCIM1228 and Δ*mig3* secretome at 10% substrate load, 3% enzyme load for 24 hours. All experiments were repeated three times; statistical significance was determined by a one-tailed, unequal variance t-test.

**Table 2.**
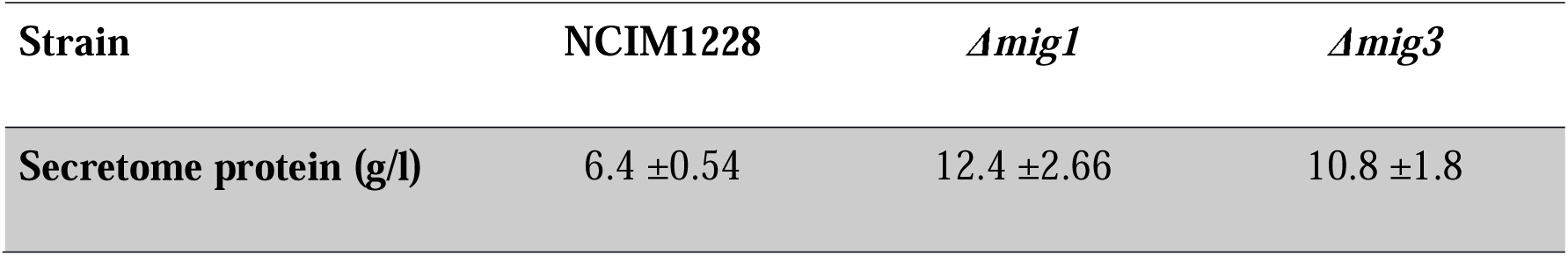
Total protein secreted by *T. pinophilus* and deletion mutants in RCM medium. . Protein content was measured by the BCA method. Values represent the mean ± standard deviation of biological data in triplicate. The significance of the data was evaluated by t-tests and was found to be <0.01.

| Strain | NCIM1228 | *Δmig1* | *Δmig3* |
| --- | --- | --- | --- |
| Secretome protein (g/l) | 6.4 ±0.54 | 12.4 ±2.66 | 10.8 ±1.8 |

### 3.3 TpMig1 and TpMig3 are expressed under both repressing and derepressing conditions

To examine the expression of native TpMig1 and TpMig3 proteins in *T. pinophilus*, we generated specific polyclonal antibodies against the zinc-finger domains of each repressor (Fig. 3a; Fig. S3). Both antibodies selectively recognized their respective recombinant antigens and did not cross-react (Fig. 3a). Using these antibodies, we detected endogenous TpMig1 and TpMig3 in whole-cell protein extracts from glucose-grown cultures. Immunoblotting with anti-Mig1 identified a single band of approximately 18 kDa corresponding to the previously described truncated TpMig1 protein (TpMig1^134^), which was absent in the Δ*mig1* strain (Fig. 3b). This confirmed that TpMig1^134^ is stably expressed *in vivo* and is likely responsible for the residual CCR observed in Δ*mig1* strains. In contrast, immunoblotting with anti-Mig3 consistently detected two bands in the parental strain: a ∼100 kDa band corresponding to full-length TpMig3 and a smaller ∼70 kDa species (Fig. 3c). Both bands were absent in Δ*mig3*, indicating that they represent Mig3-derived products. The presence of multiple TpMig3 species suggests post-translational processing or differential stability of the protein.

**Fig. 3.**
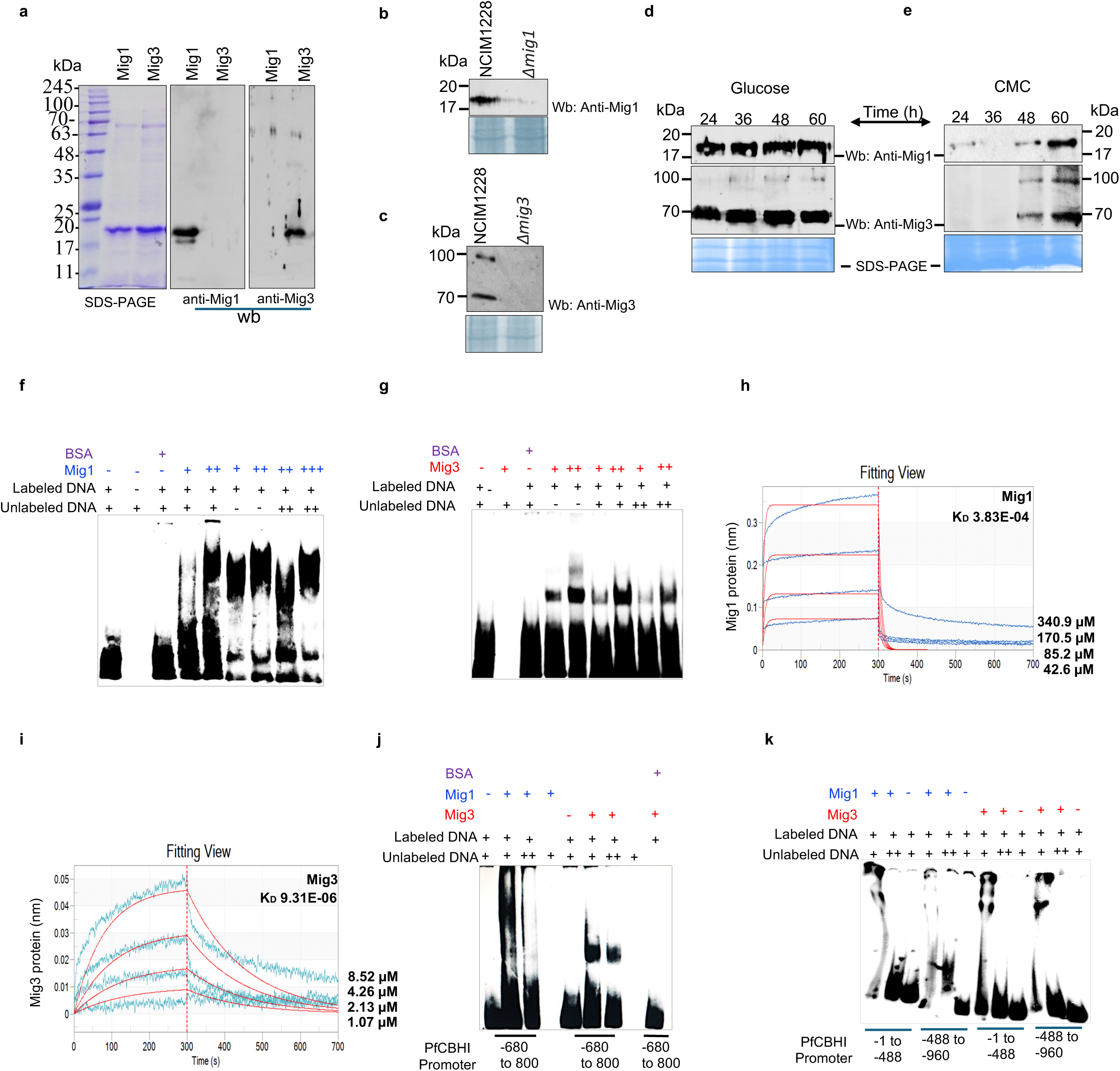
Recombinant TpMig1 and TpMig3 zinc finger domains bind to the GCGGGG DNA motif. **a,** Recombinant expression of 6X-His tagged zinc finger domains of TpMig1 and TpMig3 in *E. coli*, and western blotting of recombinant zinc fingers with anti-Mig1 and anti-Mig3 antibodies confirming the specificities of antibodies for TpMig1 or TpMig3. **b,** western blotting of whole mycelial protein extract of NCIM1228 and Δ*mig1* with anti-mig1. **c,** Anti-mig1 western blotting of whole mycelial protein extract of NCIM1228 and Δ*mig1* cultured in glucose for 36 hours. Anti-mig1 and anti-Mig3 western blotting of whole mycelial protein extract of NCIM1228 and Δ*mig3* cultured in (**d**) glucose and (**e**) carboxymethyl cellulose (CMC) for different time points. EMSA profile of recombinant (**f**) Mig1 and (**g**) Mig3 zinc finger domains to dsDNA having ‘CCATAT<u>GCGGGG</u>TTTGTTCC’ sequence from the *cbh1* promoter. BSA was taken as a negative control. BLI plot showing association and dissociation curve of (**h**) Mig1 and (**i**) Mig3 DNA binding domains to dsDNA having ‘CCATAT<u>GCGGGG</u>TTTGTTCC’ sequence from the *cbh1* promoter. **j,** EMSA profile of recombinant Mig1 and Mig3 DNA binding domains to the -680 to -800 bp region of the *cbh1* promoter. **k**, EMSA profile of Mig1 and Mig3 DNA binding domains to -1 to -488 bp, and - 488 to -960 bp regions of the *cbh1* promoter. All western blot and EMSA experiments were repeated three times.

We next examined the temporal expression profiles of TpMig1 and TpMig3 under repressing (glucose) and derepressing (cellulose) conditions. When cultured in glucose, both repressors were expressed at relatively constant levels over the entire growth period examined, with no substantial decline even after prolonged incubation (Fig. 3d). For TpMig3, the truncated form consistently accumulated to higher levels than the full-length protein. Under cellulosic conditions, neither TpMig1 nor TpMig3 was detected during early growth stages. TpMig1 expression became detectable only after prolonged incubation (60 h), coinciding with the onset of robust cellulolytic activity (Fig. 3e). Similarly, TpMig3 expression was observed only at later time points (48–60 h), at which both full-length and truncated forms accumulated (Fig. 3e). Together, these results demonstrate that TpMig1 and TpMig3 are not restricted to glucose-rich conditions but are expressed under both repressing and derepressing growth regimes, with delayed accumulation during growth on cellulose. The persistence of both repressors under derepressing conditions suggests regulatory roles beyond simple glucose-dependent repression and supports their involvement in broader metabolic coordination.

### 3.4. TpMig3 zinc finger exhibits a higher affinity GCGGGG motif than TpMig1

Mig1/CreA family repressors recognize a conserved 5′-SYGGRG-3′ motif. To determine whether TpMig3 binds this canonical element, we examined DNA binding of recombinant zinc-finger domains of TpMig1 and TpMig3 using electrophoretic mobility shift assays (EMSA). A 20-bp fragment of the *cbh1* promoter containing the GCGGGG motif located 763 bp upstream of the transcription start site was used as a probe. Both TpMig1 and TpMig3 zinc-finger domains specifically bound the GCGGGG-containing probe, as evidenced by retarded probe mobility relative to control reactions containing bovine serum albumin (Fig. 3f,g). Binding specificity was confirmed by competition with excess unlabeled probe, which restored probe mobility in a concentration-dependent manner. However, the two proteins exhibited distinct binding behaviours. TpMig3 bound the probe efficiently at lower protein concentrations and displayed a linear increase in binding with increasing protein levels. In contrast, TpMig1 required higher protein concentrations to achieve detectable binding and exhibited a steeper, non-linear binding response, suggesting cooperative interactions. To quantitatively compare DNA-binding affinities, we measured association and dissociation kinetics using biolayer interferometry. TpMig3 displayed a higher association rate constant and a lower dissociation rate constant than TpMig1, resulting in a substantially stronger apparent binding affinity for the GCGGGG motif (Fig. 3h,i). TpMig1, by contrast, showed slower association and faster dissociation kinetics, consistent with weaker and more transient DNA interactions. Gel permeation chromatography further indicated that TpMig1 exists predominantly as a dimer in solution (Fig. S4), suggesting that cooperative binding may contribute to its DNA interaction properties. We next assessed whether these binding differences persisted on larger promoter fragments. EMSA performed with three *cbh1* promoter fragments (120–500 bp) revealed that both TpMig1 and TpMig3 were capable of retarding all fragments to varying extents (Fig. 3j,k). Notably, the *cbh1* promoter has one GCGGGG and three SYGGRG motifs and both proteins showed binding to the promoter fragment having either the canonical GCGGGG or SYGGRG motifs, indicating that TpMig1 and TpMig3 can associate with both canonical GCGGGG and divergent SYGGRG regulatory elements (16). Importantly, the characteristic binding patterns observed with the short probe were retained with larger fragments, supporting intrinsic differences in DNA-binding behavior between the two repressors. Together, these results demonstrate that TpMig1 and TpMig3 recognize overlapping DNA targets but differ markedly in binding affinity and kinetics, with TpMig3 exhibiting stronger and more stable interactions with canonical CCR motifs. These differences likely contribute to their overlapping yet non-identical regulatory roles in carbon catabolite repression.

### 3.5. TpMig1 and TpMig3 genes form a synthetic lethal pair

To test whether TpMig1 and TpMig3 together constitute the core CCR machinery in *T. pinophilus*, we attempted to generate a double deletion mutant lacking both repressors. Repeated efforts to disrupt *mig3* in the Δ*mig1* background, and conversely *mig1* in Δ*mig3*, consistently failed to yield viable transformants. To exclude transformation inefficiency as a confounding factor, we next performed genetic crosses between Δ*mig1* and Δ*mig3* strains, followed by non-random spore analysis. In multiple independent mating experiments, large numbers of hygromycin- or zeocin-resistant progeny were recovered, corresponding to the respective single mutants. However, no viable progeny resistant to both antibiotics were obtained, despite screening >10,000 spores per cross (Fig. 4a; Table 3). Given that non-allelic single mutants typically yield double mutants among non-parental ditype asci, the systematic absence of Δ*mig1*Δ*mig3* progeny strongly suggests that simultaneous loss of both genes is lethal.

**Fig. 4.**
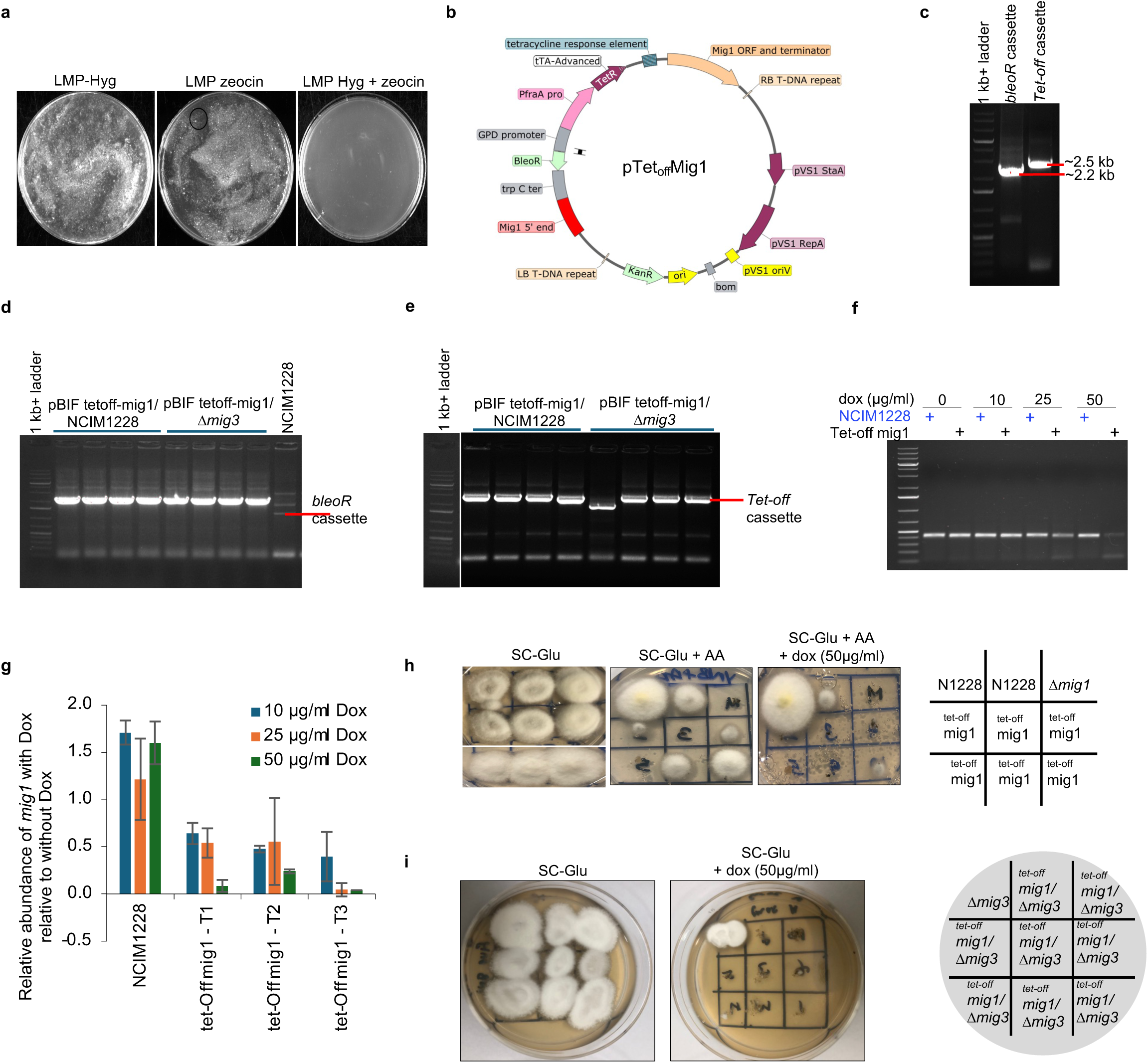
*TpMig1* and *TpMig3* form a synthetic lethal pair. **a,** Δ*mig1* and Δ*mig3* were induced for mating, and random spore analysis after mating was conducted. 10^6^ spores were plated in the presence of hygromycin, zeocin, and both hygromycin and zeocin. None of the spores survived on both hygromycin and zeocin. **b,** vector map of pTet_off_Mig1 plasmid having *TpMig1^134^* gene under Tet_off_-regulated promoter. **c,** PCR showing the amplification of the bleomycin resistance cassette and Tet_off_-regulated promoter in pTet_off_Mig1. PCR showing the amplification of (**d**) bleomycin resistance cassette and (**e**) Tet_off_-regulated promoter in pTet_off_Mig1transformants in NCIM1228 and Δ*mig3* background. **f,** pTet_off_Mig1 NCIM1228 transformants were cultured in glucose with different concentrations of doxycycline, and RT-PCR amplification of *mig1* mRNA exhibiting inhibiting effects of 10mg/ml doxycycline on *mig1* expression. **g,** qPCR showing the abundance of *mig1* transcript in the presence of doxycycline at different concentrations. **h,** growth profile of NCIM1228, Δ*mig1*, and p_tet-off_*mig1*NCIM1228 on allyl alcohol with and without doxycycline. **i,** growth profile of Δ*mig3*, and p_tet-off_*mig1*/Δ*mig3* transformants with and without doxycycline. The experiments were conducted on 3-10 integrative transformants three times.

**Table 3.**
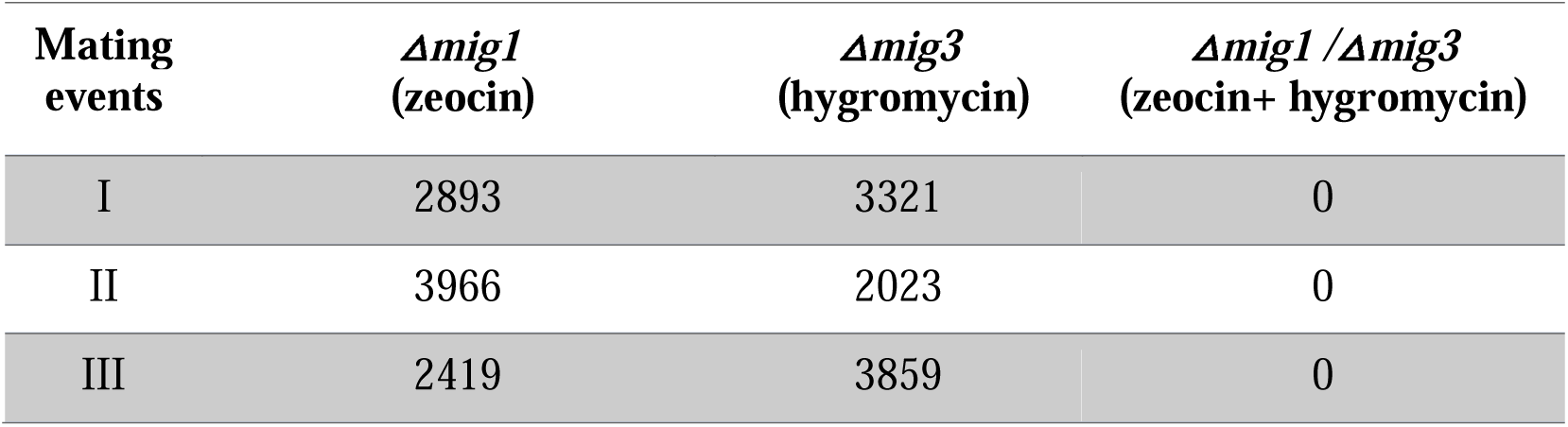
Mating events of Δ*mig1* and Δ*mig3*, followed by random spore analysis. The table shows the number of colonies formed after 10000 spores after mating events were plated in the presence of hygromycin, zeocin, and both hygromycin, and zeocin. The experiment was repeated three times.

To directly validate this synthetic lethal interaction, we employed a tetracycline-regulated conditional expression system. The Tet-Off conditional expression system utilises the tetracycline-controlled transactivator (tTA), in which tTA binding to the responsive operator sequence tetO is prevented by tetracycline/doxycycline, enabling quantitative reduction of gene expression. *mig1* was placed under control of a Tet-Off promoter, allowing doxycycline-dependent repression of *mig1* expression (Fig. 4b). The Tet-Off-*mig1* construct was introduced into both the wild-type and Δ*mig3* backgrounds, and correct genomic integration was confirmed by PCR (Fig. 4c–e). In the wild-type background, doxycycline treatment resulted in a dose-dependent reduction of *mig1* transcript levels, as verified by semi-quantitative and quantitative RT-PCR, with 50 μg/ml doxycycline sufficient to achieve strong repression after 48 h (Fig. 4f,g). Functional downregulation of *mig1* was further confirmed by the acquisition of allyl alcohol sensitivity under repressing conditions, consistent with impaired CCR (Fig. 4h). Strikingly, repression of *mig1* expression in the Δ*mig3* background resulted in complete growth arrest on glucose-containing medium in the presence of doxycycline, whereas growth was unaffected in the absence of doxycycline (Fig. 4i). This conditional lethality unequivocally demonstrates that *mig1* and *mig3* constitute a synthetic lethal gene pair, and that expression of at least one of the two catabolite repressors is essential for viability in *T. pinophilus*. Together, these genetic analyses reveal an unexpected essential function for fungal catabolite repressors that extends beyond carbon repression and suggests that TpMig1 and TpMig3 jointly sustain core cellular processes required for survival.

### 3.6 Transcriptomic profiling reveals redundant control of mitochondrial and ribosomal biogenesis by TpMig1 and TpMig3

The synthetic lethality observed between *mig1* and *mig3* suggested that these catabolite repressors may redundantly regulate a set of essential genes required for cellular viability. To identify such shared targets and pathways, we performed RNA-seq–based transcriptomic profiling of log-phase cultures of the wild-type strain NCIM1228, Δ*mig1*, and Δ*mig3* grown under repressing (glucose) and de-repressing (cellulose) conditions. Hierarchical clustering of global expression profiles revealed a clear separation between glucose- and cellulose-grown transcriptomes (p < 0.05), with high reproducibility among biological replicates (Fig. 5a). Principal component analysis (PCA) further confirmed carbon source as the dominant driver of transcriptional variation (Fig. 5b). While glucose-grown samples clustered closely when all strains were considered together, PCA restricted to glucose-grown transcriptomes resolved distinct strain-specific clusters, indicating subtle but reproducible transcriptional differences arising from *mig1* or *mig3* deletion even under repressing conditions (Fig. 5c). Comparative differential expression analysis identified 341, 761, and 634 genes upregulated, and 201, 416, and 258 genes downregulated in cellulose relative to glucose in NCIM1228, Δ*mig1*, and Δ*mig3*, respectively. Clustering of differentially expressed genes revealed a substantial core cellulose-responsive regulon shared across all three strains (Fig. 5d). Genes commonly upregulated in cellulose primarily encoded (i) carbohydrate-active enzymes involved in cell wall degradation, (ii) meiotic proteins, and (iii) members of the major facilitator superfamily (MFS) of transporters (Table 4). Notably, both the magnitude of induction and the number of genes within these pathways followed the trend NCIM1228 > Δ*mig3* > Δ*mig1*, consistent with progressive derepression of carbon-responsive pathways in the mutants. Conversely, genes commonly downregulated in cellulose across all strains were enriched for metabolic biosynthesis pathways, including amino acid, organic acid, sterol, nucleobase, and secondary metabolite biosynthesis, as well as galactose, starch, and sucrose metabolism (Table 4). However, the extent of repression within these pathways differed markedly among strains, with Δ*mig3* exhibiting the strongest reduction in genes involved in core biosynthetic processes. Strain-specific analysis further revealed that Δ*mig1* and Δ*mig3* shared distinctive transcriptional signatures not observed in the wild-type strain. Both mutants exhibited exclusive upregulation of genes encoding ABC transporters and cellulolytic enzymes in cellulose, while Δ*mig1* uniquely induced peptide transporters and Δ*mig3* selectively upregulated secondary metabolism–associated genes (Fig. 5f,g). In contrast, NCIM1228 showed exclusive enrichment of genes involved in cell signaling and cell wall organization (Fig. 5e). Strikingly, analysis of genes exclusively downregulated in the mutant strains uncovered a profound repression of genes encoding intracellular non-membrane–bound organelles, specifically ribosomes and mitochondria (Fig. 6). In Δ*mig1*, over 70 genes encoding ribosomal proteins from both large and small subunits were significantly downregulated under cellulose conditions, accompanied by repression of cap-dependent translation initiation and tRNA aminoacylation pathways (Fig. 6a–c). In parallel, more than 90 genes associated with mitochondrial structure and function were repressed, including components of the mitochondrial membrane, transcription and translation machinery, tRNA aminoacylation enzymes, RNA metabolism factors, ubiquinone biosynthesis enzymes, and the pyruvate dehydrogenase complex (Fig. 6d). A similar but quantitatively attenuated repression of ribosomal and mitochondrial gene sets was observed in Δ*mig3*, with the exception of tRNA aminoacylation, which was largely unaffected (Fig. 6e,f). Importantly, no such repression of ribosomal or mitochondrial pathways was detected in the wild-type strain, indicating that loss of either catabolite repressor compromises expression of core cellular machinery under de-repressing conditions. Together, these transcriptomic analyses reveal that TpMig1 and TpMig3 redundantly sustain mitochondrial and ribosomal biogenesis when cells transition away from glucose. The simultaneous loss of both repressors is therefore predicted to collapse central metabolic and translational capacity, providing a mechanistic explanation for the synthetic lethality observed upon dual inactivation of *mig1* and *mig3*.

**Fig. 5.**
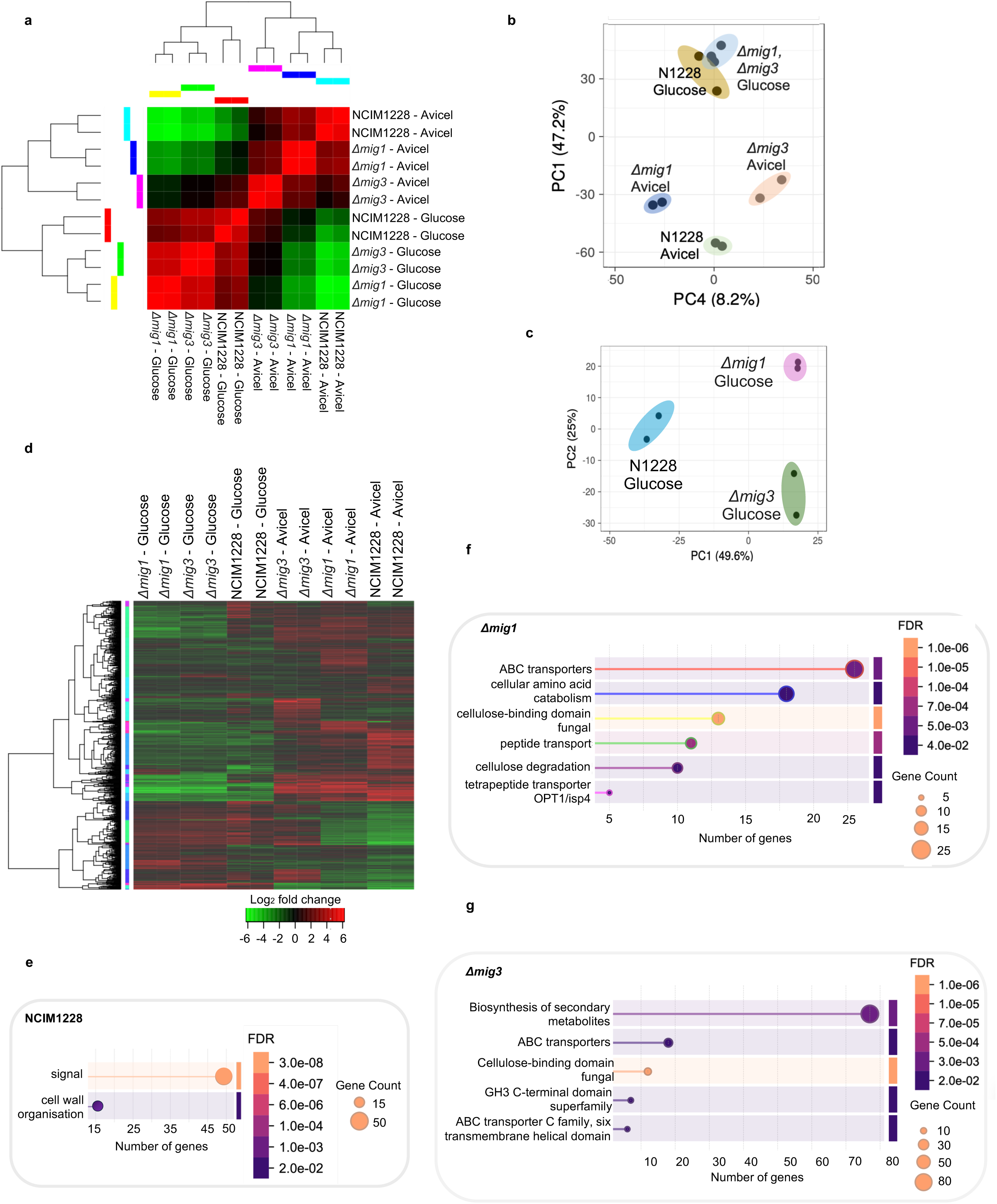
Transcriptomic profiling of NCIM1228, Δ***mig1*, and** Δ***mig3* under repressing and de-repressing conditions. a,** sample correlation matrix of NCIM1228, Δ*mig1*, and Δ*mig3* when grown in glucose and Avicel. **b,** Principal component analysis of whole transcriptome samples of NCIM1228, Δ*mig1*, and Δ*mig3* cultured in glucose and Avicel. **c,** Principal component analysis of whole transcriptome samples of NCIM1228, Δ*mig1*, and Δ*mig3* cultured in glucose. **d,** the heatmap representing hierarchical clustering of normalized FPKM abundance (log_2_) of differentially expressed genes of NCIM1228, Δ*mig1*, and Δ*mig3*. Differentially expressed genes were analyzed by STRING to identify pathways represented by them. **e,** differentially upregulated pathways in cellulose cultures of NCIM1228 compared to glucose cultures. **f**, differentially upregulated pathways in cellulose cultures of Δ*mig1* compared to glucose cultures. **g**, differentially upregulated pathways in cellulose cultures of Δ*mig1* compared to glucose cultures.

**Fig. 6.**
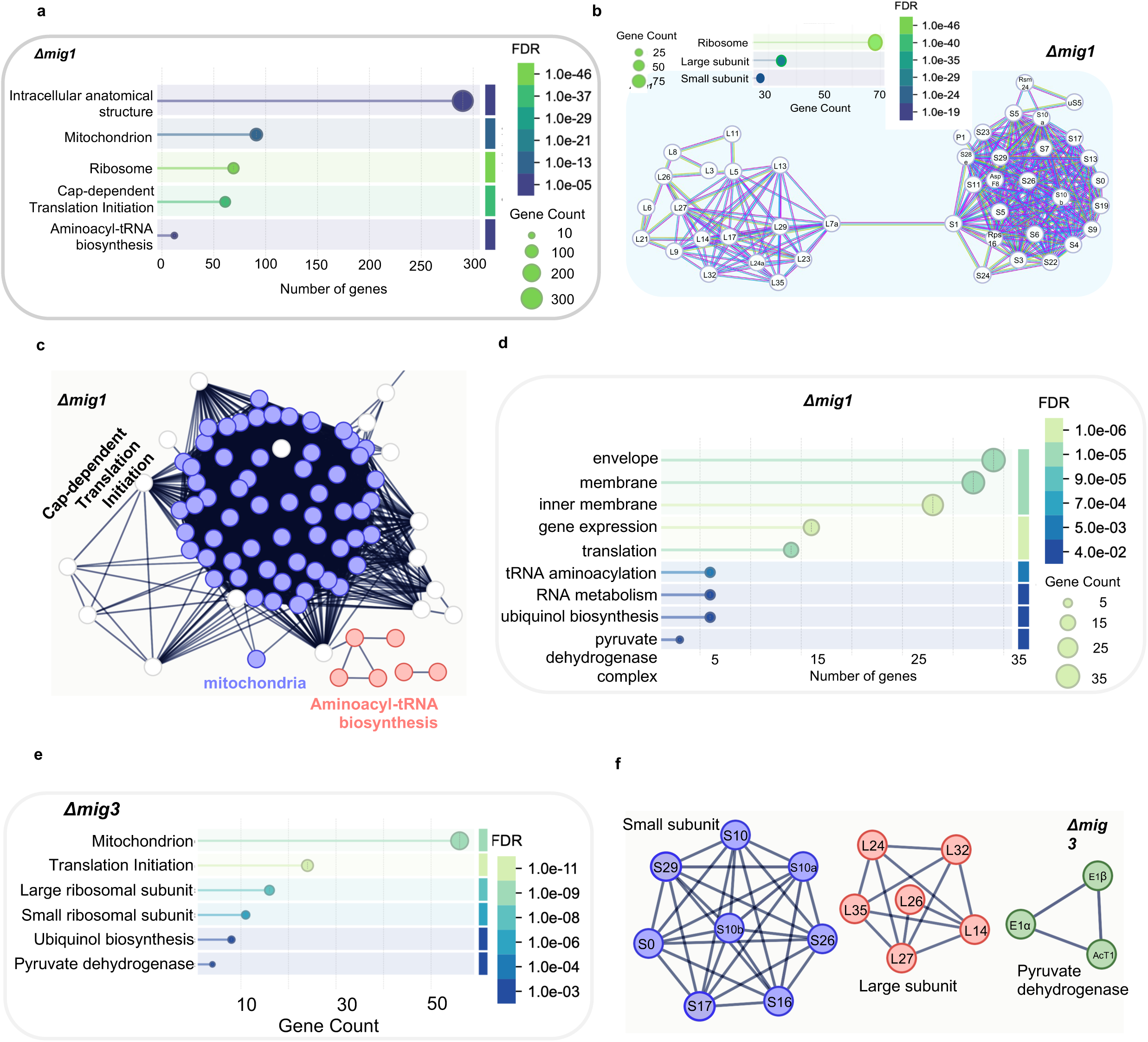
Pathways downregulated in Δ*mig1* and Δ*mig3* under de-repressing conditions relative to their transcriptome in glucose. Differentially expressed genes were analysed by STRING to identify pathways represented by them. **a,** Major pathways represented by Δ*mig1* differentially downregulated cellulose transcriptome relative to glucose. **b,** down-regulated genes in Δ*mig1* cellulose transcriptome representing the large and small subunit of ribosomes. **c,** down-regulated genes in Δ*mig1* cellulose transcriptome encoding for mitochondrial components and function, Cap-dependent translation initiation, and Amino-acyl tRNA biosynthesis. **d,** Major mitochondrial pathways represented by down-regulated genes in Δ*mig1* cellulose transcriptome. **e,** Major pathways represented by Δ*mig3* differentially downregulated transcriptome. **f,** down-regulated genes in Δ*mig1* glucose transcriptome representing large and small subunit of ribosomes, and pyruvate dehydrogenase complex.

**Table 4.** Common pathways enriched by the genes found upregulated and downregulated in cellulose compared to glucose.

| Common upregulated pathways in cellulose | Number of genes enriched |  |  | Total genes representing the pathway |
| --- | --- | --- | --- | --- |
| | N1228 | $\Delta mig1$ | $\Delta mig3$ | |
| Xylan degradation/ β-glucosidase/ cell wall degradation | 12 | 26 | 24 | 68 |
| Meiosis | 17 | 28 | 23 | 216 |
| MFS transporter superfamily | 35 | 64 | 52 | 477 |
| I. Inositol transporters | 16 | 27 | 22 | 123 |
| II. Sugar transporters | 11 | 20 | 16 | 73 |
| III. Transporters of organic acids, metal ions, and amines | 17 | 30 | 24 | 193 |
| **Common downregulated pathways in cellulose** |  |  |  |  |
| Biosynthesis of organic acids | 34 | 55 | **19** | 367 |
| Biosynthesis of amino acids | 32 | 42 | **11** | 282 |
| Biosynthesis of secondary metabolites | 61 | 91 | 57 | 1092 |
| Starch and sucrose metabolism, and Galactose metabolic process | 8 | 14 | 12 | 73 |
| Sterol biosynthetic process | 6 | - | 9 | 50 |
| Nucleobase biosynthetic process | 4 | 5 | 7 | 18 |

### 3.7 TpMig1 and TpMig3 act as transcriptional activators of oxidative phosphorylation genes under repressing conditions

To determine whether TpMig1 and TpMig3 influence gene expression under carbon-repressing conditions, we compared the glucose-grown transcriptomes of Δ*mig1* and Δ*mig3* with that of the wild-type strain NCIM1228. Of the 10,843 transcripts detected under glucose conditions, deletion of *mig1* altered the expression of 167 genes, including 110 downregulated and 57 upregulated genes (Fig. 7a). Similarly, deletion of *mig3* affected the expression of 108 genes, with 68 genes downregulated and 40 genes upregulated relative to the wild type (Fig. 7b). To infer the regulatory roles of TpMig1 and TpMig3, differentially expressed genes were grouped based on their direction of regulation in the mutant strains. This analysis identified gene sets whose expression was reduced in Δ*mig1* or Δ*mig3*, indicating transcriptional activation by TpMig1 or TpMig3, respectively, as well as gene sets showing increased expression, consistent with classical repression (Fig. 7c). Importantly, a distinct subset of genes was downregulated in both Δ*mig1* and Δ*mig3*, suggesting redundant activation by the two repressors. Expression patterns across these gene sets were validated by quantitative PCR analysis of 55 representative genes (Fig. 7d–f). Gene set enrichment analysis (GSEA) of the combined differentially expressed genes revealed a striking functional dichotomy. Genes upregulated in the mutant strains were enriched for transporters and metabolic pathways involved in sugar, glucoside, polyol, and carboxylate utilization, consistent with relief from carbon catabolite repression (Fig. 7g). In contrast, genes downregulated in Δ*mig1* and Δ*mig3* were strongly enriched for pathways associated with mitochondrial electron transport, including NAD/NADP- and FAD/FMN-binding proteins, ubiquinone and ubiquinol biosynthesis, and oxidative phosphorylation (Fig. 7g). We next focused on genes whose expression was reduced in both Δ*mig1* and Δ*mig3*, as these represent potential shared essential targets underlying the observed synthetic lethality. Manual curation of this redundantly activated gene set revealed a strong enrichment for genes encoding oxidoreductases involved in oxidative phosphorylation, followed by smaller groups of transcription factors, kinases, methyltransferases, carbohydrate-active enzymes, and MFS transporters (Fig. 7h). Consistently, GSEA identified oxidoreductase activity—particularly enzymes acting on paired donors with incorporation or reduction of molecular oxygen—as the most significantly enriched functional category within this gene set (Fig. 7i). Together, these results demonstrate that TpMig1 and TpMig3 function not only as repressors of alternative carbon utilization pathways, but also as transcriptional activators of genes required for mitochondrial respiration, even under glucose-repressing conditions. The redundant activation of oxidative phosphorylation genes by TpMig1 and TpMig3 provides a mechanistic explanation for the synthetic lethality observed upon their simultaneous inactivation and reveals an unexpected role for catabolite repressors in sustaining respiratory metabolism.

**Fig. 7.**
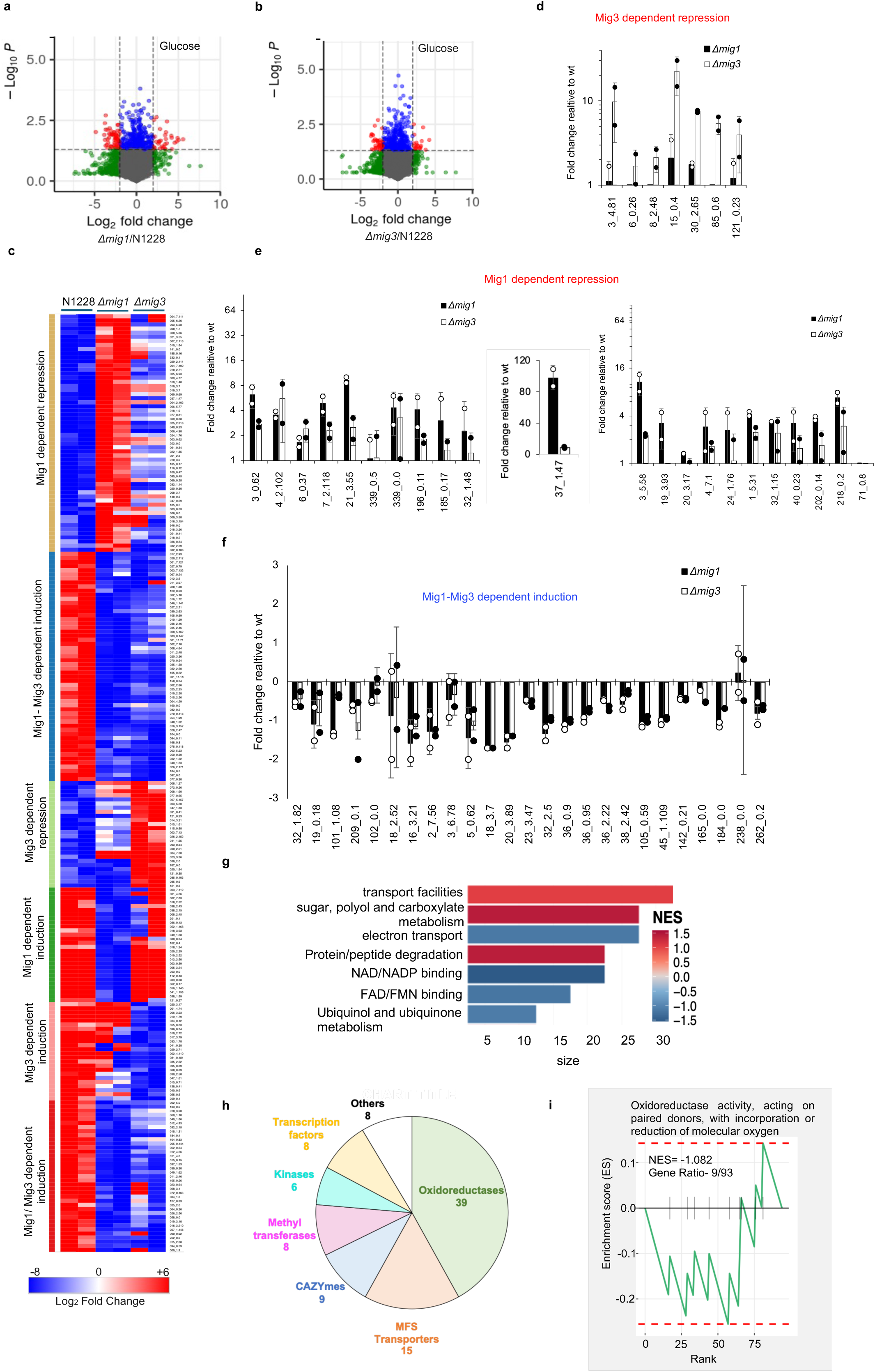
Common Pathways downregulated in Δ*mig1*, and Δ*mig3* relative to NCIM1228 under repressing conditions. **a,** volcano plot of differentially expressed genes in Δ*mig1* with respect to NCIM1228 in glucose. **b,** volcano plot of differentially expressed genes in Δ*mig3* with respect to NCIM1228 in glucose. **c,** heatmap exhibiting hierarchical clustering of normalized FPKM abundance (log_2_) of common downregulated gene set of Δ*mig1* and Δ*mig3* glucose transcriptome. **d-f,** quantitative RT-PCR of selected genes representing different clusters in NCIM1228, Δmig1 and Δmig3 grown in MM glucose for 48 h. The C(T) value in Δ*mig1* and Δ*mig3* was normalized to that in NCIM1228. **g,** gene set enrichment analysis of upregulated and downregulated common gene cluster of Δ*mig1* and Δ*mig3* with respect to NCIM1228 (NES- Normalized Enrichment Score). **h,** Manual clustering of downregulated common gene set based on their function. **i,** gene set enrichment analysis of common downregulated gene cluster of Δ*mig1* and Δ*mig3* with respect to NCIM1228.

### 3.8 ChIP-seq reveals direct binding of TpMig1 and TpMig3 to promoters of genes involved in oxidative phosphorylation

Transcriptomic analyses identified oxidative phosphorylation, mitochondrial function, and ribosomal biogenesis as major pathways redundantly downregulated in Δ*mig1* and Δ*mig3* strains, suggesting that TpMig1 and TpMig3 may directly activate genes essential for respiratory metabolism. To test this hypothesis and distinguish direct from indirect regulatory effects, we performed genome-wide ChIP-seq analysis of TpMig1 and TpMig3 in the wild-type strain NCIM1228 under carbon-repressing conditions. ChIP-seq experiments were carried out using anti-Mig1 and anti-Mig3 antibodies on NCIM1228 cultures grown in 4% glucose for 48 h. After normalization to input controls, we identified binding peaks proximal to a total of 295 genes in TpMig1 and 270 in TpMig3ChIP-seq data (Fig. 8a). Heatmap analysis revealed strong enrichment of both TpMig1 and TpMig3 binding in regions proximal to transcription start sites (TSSs), with the highest peak densities observed within 1 kb upstream of annotated genes (Fig. 8a, b). Consistent with this observation, 61.7% of TpMig1 peaks and 67.4% of TpMig3 peaks were located within the ≤1 kb promoter region, whereas smaller fractions were detected in the 1–2 kb, 2–3 kb, or distal intergenic regions (Fig. 8c). Comparison of TpMig1 and TpMig3 ChIP-seq datasets revealed substantial overlap between the two regulons. A total of 122 TpMig1 peaks and 103 TpMig3 peaks overlapped, corresponding to 93 shared target genes, with co-occupancy predominantly centered around the TSS (Fig. 8d). This overlap supports a model in which TpMig1 and TpMig3 function as co-regulators at a subset of promoters, consistent with their genetic redundancy and synthetic lethal interaction. De novo motif discovery of co-bound regions identified enrichment of the canonical Cre-like motif (5′-GCGGGG-3′), consistent with direct DNA binding by both factors (Fig. 8e, f). Additionally, regulatory elements representing GC-rich Mig1 variant motif, SYGGRG, stress-responsive element (STRE; GGGG/CCCC), and G-box elements were also highly enriched, suggesting that TpMig1 and TpMig3 bind promoters with complex regulatory architectures (Fig. 8g). In addition, motif enrichment analysis revealed binding sites for several transcription factors, including Hap4, Sfp1, Rlr1, Xbp1, Nhp10, Put3, and Aft2 (Fig. 8h), indicating potential cooperative or hierarchical regulation of shared target genes.

**Fig. 8.**
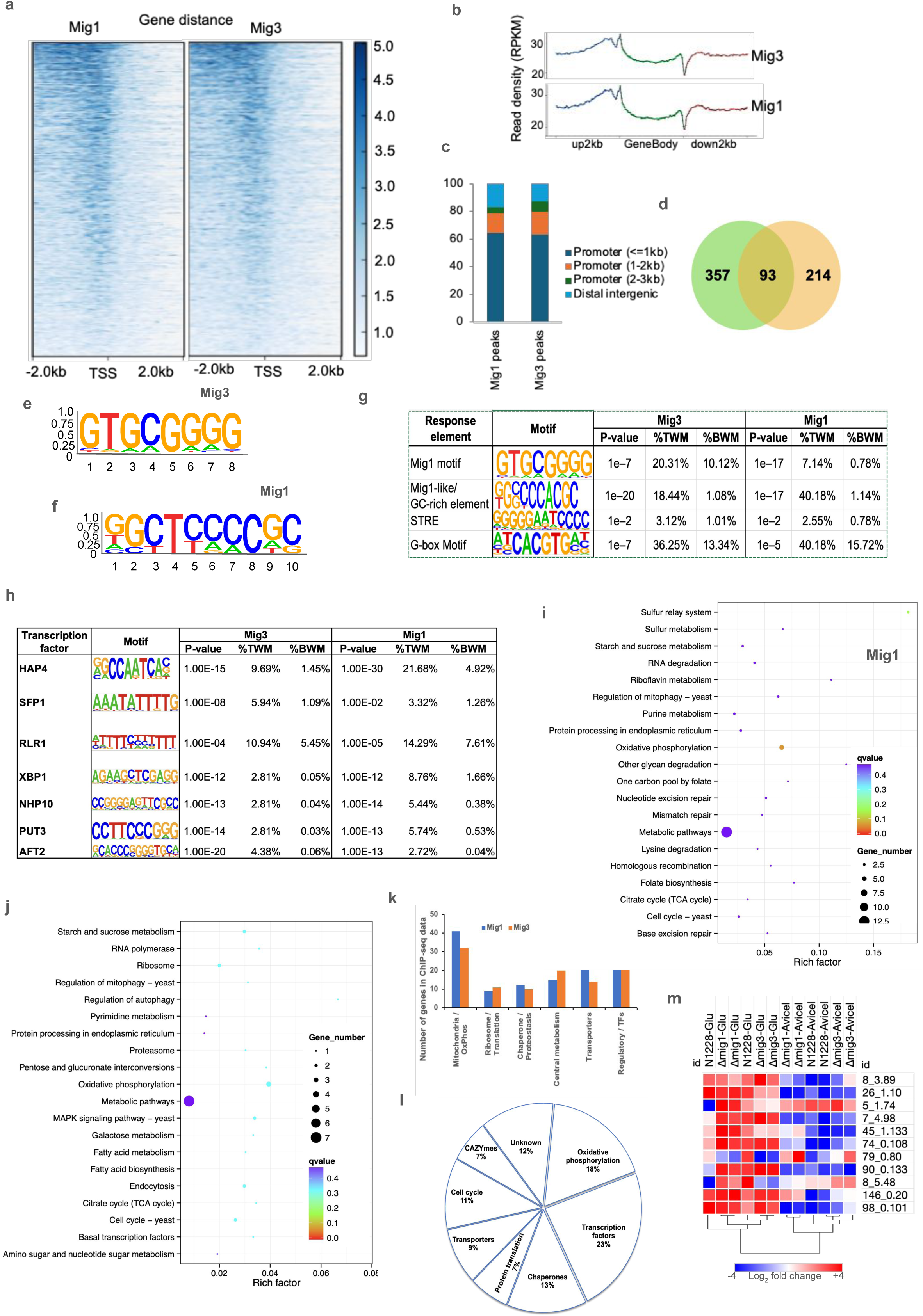
Genome-wide analysis of TpMig1 and TpMig3 DNA binding sites by ChIP-seq. **a,** heatmap exhibiting DNA binding of TpMig1 and TpMig3. **b,** the distribution of reads on the genome. **c,** percentage of TpMig1 and TpMig3 DNA binding peaks with respect to the transcription start site. **d,** Venn diagram showing overlapping peaks among Mig1 and Mig3 ChIP-seq data. GCGGGG DNA-binding motif identified in **(e)** Mig3 and **(f)** Mig1 ChIP-seq data. **g,** Percentage of Variants of SYGGRG DNA-binding motifs identified in TpMig1 and TpMig3 ChIP-seq data. **h,** DNA-binding motifs identified in TpMig1 and TpMig3 ChIP-seq data, which are also binding sites for other conserved eukaryotic transcription factors. KEGG enrichment scatter plot of **(i)** TpMig1, and **(j)** TpMig3. **k,** Manual gene function-based clustering of gene sets being regulated by both TpMig1 and TpMig3. **l,** gene function-based clustering of the common gene set identified in both TpMig1 and TpMig3 ChIP-seq data. **m,** heatmap exhibiting comparative expression profiling co-bound genes encoding for oxidative phosphorylation genes in NCIM1228, Δ*mig1* and Δ*mig3* cultured in glucose and Avicel. Hierarchical clustering divided the samples into two major groups.

Gene set enrichment analysis of TpMig1- and TpMig3-bound genes showed strong enrichment for metabolic pathways, with oxidative phosphorylation emerging as one of the most significantly overrepresented functional categories (Fig. 8i, j). Detailed functional annotation revealed strong enrichment for genes of mitochondrial and oxidative phosphorylation genes, including multiple components of the electron transport chain and ATP synthesis machinery (Fig. 8k). Strikingly, both datasets also showed substantial over-representation of ribosomal proteins, translation factors, and rRNA/tRNA processing enzymes, together with numerous molecular chaperones and proteostasis regulators. Additional categories included central carbon metabolism enzymes and membrane transporters (Fig. 8k). Next, analysis of the 93 co-bound genes also revealed predominance of transcriptional regulators (23%), enzymes of oxidative phosphorylation (18%), molecular chaperones (13%), cell cycle regulators (11%), carbohydrate-active enzymes (9%), and transporters (7%) (Fig. 8l). Furthermore, genes related to oxidative phosphorylation code for proteins involved in iron sequestration and components of complex I, complex II, complex III, and complex IV of electron transport chain (Table 5). Notably, the majority of the co-bound oxidative phosphorylation targets were downregulated in cellulose (Fig. 8m).

**Table 5.** ChIP-seq data showing binding sites (BS) of TpMig1 and TpMig3 from the transcriptional start site (TSS) of co-regulated genes of oxidative phosphorylation. The genes were also downregulated in cellulose in NCIM1228, Δ*mig1* and Δ*mig3*.

| Gene ID | Gene Annotation | Gene Function | DNA strand | Mig1 BS from TSS | DNA strand | Mig3 BS from TSS |
| --- | --- | --- | --- | --- | --- | --- |
| 8_3.89 | COX13-cytochrome c oxidase subunit 6A | last step of ETC | + | 0 | - | 0 |
| 26_1.10 | ARH1- NADPH ubiquinone reductase | Essential, FE-S and iron homeostasis | + | 23 | + | 0 |
| 5_1.74 | QFR Quinol-fumarate reductase | complex II of ETC, accepts quinol | + | 0 | - | 0 |
| 7_4.98 | QCR6-ubiquinol-cytochrome-c reductase subunit | complex III of ETC | - | 49 | - | 0 |
| 45_1.133 | FAD+-membrane bound oxidoreductase | complex II of ETC | + | -488 | - | -508 |
| 74_0.108 | cytochrome c oxidase subunit | last step of ETC | - | -435 | - | -437 |
| 167_0.1 | Fo complex subunit of F-type ATP synthase | ATP synthesis | - | -131 | + | -430 |
| 79_0.8 | FRE2- Ferric/Cupric reductase | Iron import | - | -1079 | - | -1109 |
| 8_5.48 | Mix17-mitochondrial intermembrane space protein | oxidative phosphorylation | + | 0 | - | 43 |

Taken together, these data demonstrate that TpMig1 and TpMig3 directly bind the promoters of genes involved in oxidative phosphorylation and mitochondrial metabolism under glucose-repressing conditions. Combined with transcriptomic evidence showing reduced expression of these genes under repressing conditions, the ChIP-seq results establish TpMig1 and TpMig3 as direct transcriptional activators of essential respiratory pathways. This direct regulatory role provides a mechanistic explanation for the synthetic lethality observed upon simultaneous loss of TpMig1 and TpMig3 and reveals an unexpected coupling of carbon catabolite repression with mitochondrial gene regulation.

### 3.9 Deletion of mig1 or mig3 increases sensitivity to inhibitors of oxidative phosphorylation and ribosome biogenesis

Transcriptomic and ChIP-seq analyses showed that TpMig1 and TpMig3 directly activate genes involved in oxidative phosphorylation and ribosome biogenesis under both repressing and derepressing conditions. Because simultaneous loss of both repressors is synthetically lethal, we hypothesized that deletion of either gene compromises essential mitochondrial and translational functions, leading to increased sensitivity to pathway-specific inhibitors. To test this, we assessed the sensitivity of Δ*mig1* and Δ*mig3* strains to antifungal compounds targeting sterol biosynthesis, rRNA/tRNA synthesis, and oxidative phosphorylation. Epoxiconazole (sterol biosynthesis), Metalaxyl (RNA polymerase I), and six mitochondrial inhibitors targeting complexes I–III, alternative oxidase, and ATP coupling were tested, with Carbendazim (microtubule inhibitor) included as a control (Fig. 9a). At EC□□ concentrations determined for the wild-type strain, Δ*mig1* and Δ*mig3* showed growth comparable to NCIM1228 in the presence of Epoxiconazole, SHAM, TTFA, and Pyraclostrobin (Fig. 9b). In contrast, both mutants exhibited moderate sensitivity to Metalaxyl, indicating impaired ribosome biogenesis. Notably, Cyazofamid caused severe growth inhibition in both deletion strains, revealing pronounced vulnerability of mitochondrial complex III. Fluazinam treatment resulted in moderate growth defects in Δ*mig1* and severe defects in Δ*mig3*, suggesting differential contributions of the two repressors to maintaining mitochondrial ATP synthesis. No increased sensitivity was observed with Carbendazim. These results functionally validate the transcriptomic and ChIP-seq data and demonstrate that deletion of either *mig1* or *mig3* compromises cellular tolerance to perturbations in oxidative phosphorylation and ribosome biogenesis, providing a mechanistic basis for their synthetic lethal interaction.

**Fig. 9.**
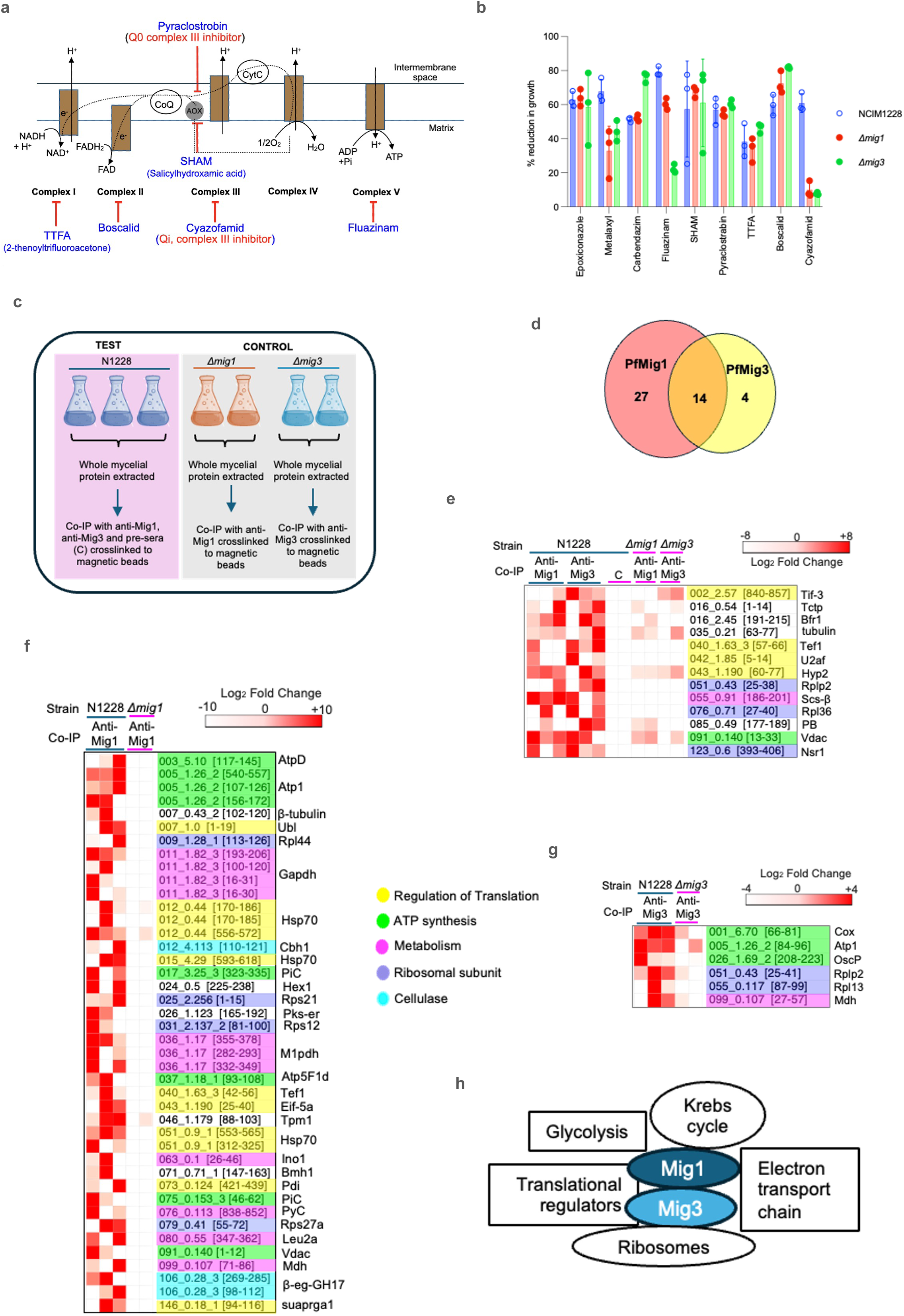
Synthetic lethality of double deletion of TpMig1 and TpMig3 is attributed to their role in ribosomal and mitochondrial gene expression. **a,** list of fungicides targeting oxidative phosphorylation used in the study and their drug targets. **b,** deletion sensitivity profiling of Δ*mig1* and Δ*mig3* with respect to NCIM1228 in Mandel’s media having 4% glucose. The experiment was repeated three times. **c,** schematic of Co-IP LC-MS/MS to identify interacting partners of Mig1 and Mig3. **d,** venn diagram showing total interacting partners identified by Co-IP LC-MS/MS. **e,** heatmap showing normalized abundance (Log_2_) of common interacting partners of Mig1 and Mig3. **f,** heatmap showing normalized abundance (Log_2_) of interacting partners of Mig1. **g,** schematic showing the pathways represented by interacting partners of TpMig1 and TpMig3.

### 3.10 TpMig1 and TpMig3 physically associate with mitochondrial and ribosomal proteins

To investigate whether TpMig1 and TpMig3 also function at post-transcriptional levels, we characterized their protein interactomes by affinity pull-down followed by LC–MS/MS. Whole-cell extracts from glucose-grown NCIM1228 cultures were incubated with anti-Mig1 or anti-Mig3 antibodies, with pre-immune sera as controls. Across all samples, 340 proteins were identified at a 1% false discovery rate. After stringent filtering, 41 proteins were enriched in the TpMig1 interactome and 18 in the TpMig3 interactome (>2-fold over control), with 14 proteins shared between the two. Functional categorization revealed enrichment for enzymes of central carbon metabolism, ribosomal proteins and translation-associated factors, regulators of translational control, and components of oxidative phosphorylation. TpMig1 additionally interacted with the cellulolytic enzymes Cellobiohydrolase 1 and Endoglucanase GH17. Together with the transcriptional and phenotypic data, these interactions indicate that TpMig1 and TpMig3 integrate transcriptional regulation of mitochondrial and ribosomal genes with physical association to metabolic and translational machinery, reinforcing their essential role in coupling carbon sensing to energy production and protein synthesis.

## 4. Discussion

Understanding global regulatory mechanisms in filamentous fungi is critical for transforming these organisms into efficient biofactories. Carbon catabolite repression (CCR) represents one such global regulatory system that connects carbon metabolism to diverse cellular processes. Although CCR has been extensively characterized in *S. cerevisiae* and, to a lesser extent, in filamentous fungi such as *N. crassa* and *A. nidulans*, its organization in hypercellulolytic fungi remains poorly understood (27–29). In this study, we identify a previously uncharacterized C2H2 zinc finger transcription factor, TpMig3, in *T. pinophilus* that appears restricted to the order Eurotiales and functions redundantly with the canonical Cre homolog TpMig1. Disruption of *Tpmig3* resulted in phenotypes closely resembling those of Δ*mig1*, indicating that TpMig3 acts as a catabolite repressor. Despite this redundancy, repeated attempts to generate a Δ*mig1*Δ*mig3* double mutant were unsuccessful, and synthetic lethality was confirmed using a Tet-Off conditional expression system. To our knowledge, this represents the first demonstration of synthetic lethality between catabolite repressors in filamentous fungi, indicating that at least one CCR regulator is required to sustain essential cellular functions in *T. pinophilus*. TpMig1 and TpMig3 exhibit distinct biochemical properties. TpMig1 exists as a truncated ∼18 kDa protein in vivo, whereas TpMig3 is detected as both a ∼100 kDa full-length protein and a ∼70 kDa truncated form. The relative abundance of these forms under glucose and cellulolytic conditions suggests regulated processing of TpMig3, although the functional significance of this processing remains to be established. Delayed accumulation of both repressors under cellulolytic conditions further suggests that they may participate in late-stage metabolic remodeling rather than acting solely as early glucose sensors, similar to CreA regulation observed in *A. nidulans* (30). Structural and biochemical analyses revealed divergent DNA-binding behaviour between the two repressors. The zinc finger domain of TpMig1 differs from that of ScMig1 by the presence of a flexible loop, whereas TpMig3 closely resembles ScMig1. Although both proteins bind the conserved GCGGGG motif, TpMig3 binds as a monomer with measurable affinity at low concentrations, while TpMig1 binds cooperatively as a dimer. Such dimeric binding by a Cre homolog has not been reported previously and suggests distinct modes of transcriptional regulation. We hypothesize that these differences allow TpMig1 and TpMig3 to act cooperatively, providing both responsiveness and stability to CCR-mediated regulation. Synthetic lethality prompted us to identify essential pathways redundantly regulated by TpMig1 and TpMig3. Integration of transcriptomic and ChIP-seq analyses revealed that both repressors regulate genes involved in mitochondrial function and ribosome biogenesis. Although ChIP-seq detected only a subset of predicted binding sites—likely reflecting technical limitations and condition specificity—the functional 5′-GCGGGG-3′ motif identified in *T. pinophilus* is consistent with findings in *A. nidulans* (29). Comparative transcriptomics further indicated that simultaneous loss of both repressors would severely compromise the expression of genes encoding ribosomal and mitochondrial components. Downregulation of ribosomal and oxidative phosphorylation genes upon deletion of Cre homologs has been reported in *A. nidulans*, *A. niger*, and *T. reesei* (28, 29, 31), but not in *N. crassa* (27), highlighting lineage-specific differences in CCR architecture. Notably, deletion of *TpMig1* alone increased growth in *T. pinophilus*, in contrast to the growth defects observed in other fungi lacking Cre homologs (32, 33). We propose that the presence of TpMig3 buffers essential gene expression programs, preventing loss of mitochondrial and ribosomal functions while permitting increased sugar uptake and growth (16). These findings contrast with the diauxic shift in *S. cerevisiae*, where Hap4 activates respiratory genes under non-fermentative conditions (34). A Hap4 homolog is absent in *T. pinophilus* (17), consistent with its oxidative lifestyle. Our ChIP-seq data revealed overlap between TpMig1/TpMig3 binding sites and chromatin regions regulated by Hap4 in yeast, suggesting that CCR regulators in *T. pinophilus* may fulfil analogous regulatory roles. Besides Hap4, TpMig1/TpMig3 co-occupancy at promoter regions that are also bound by regulators of ribosome biogenesis (Sfp1), chromatin remodeling (Nhp10/INO80), stress/quiescence (Xbp1, STRE elements), nutrient-responsive metabolic activators (Put3), and iron homeostasis (Aft2) indicates that the two catabolite repressors are embedded in a multi-factor regulatory hub (35, 35–40). This hub integrates carbon sensing and repression with transcriptional programs for mitochondrial biogenesis, ribosomal production and stress/adaptive metabolism, thereby explaining how CCR do not merely restrain peripheral carbon-use genes but occupies central nodes of bioenergetic and translational control. Beyond transcriptional control, both repressors physically interact with proteins involved in translation, mitochondrial function, and central metabolism. These interactions suggest coordination between transcriptional and post-transcriptional regulation. We hypothesize that interactions with mitochondrial chaperones such as Hsp78 help maintain mitochondrial function during carbon-replete growth (41), while interactions with Hsp70 homologs and 14-3-3 proteins may influence Snf1/AMPK signaling (42). Additionally, interactions with glyceraldehyde-3-phosphate dehydrogenase raise the possibility that CCR regulators influence rDNA chromatin organization via Sir2-dependent mechanisms (43), linking carbon availability to ribosome biogenesis.

Based on these observations, we propose a model in which TpMig1 and TpMig3 function as metabolic gatekeepers that couple extracellular carbon availability to intracellular investment in mitochondrial and ribosomal biogenesis. We propose that under glucose-rich conditions, TpMig1 and TpMig3 promote oxidative metabolism and biomass accumulation, whereas carbon limitation triggers their degradation, reduced investment in mitochondria and ribosomes, and induction of CAZyme expression. Both TpMig1 and TpMig3 do so by activating genes required for oxidative phosphorylation and translation under glucose-rich conditions, while physically associating with components of these pathways. As carbon becomes limiting, regulated degradation of TpMig1 and TpMig3 leads to reduced anabolic investment and induction of secretory enzymes and transporters, enabling efficient utilization of alternative carbon sources.

## 5. Conclusions

In summary, this study reveals an unexpected essential role for CCR regulators in maintaining cellular integrity in filamentous fungi. By uncovering a synthetic lethal relationship between TpMig1 and the lineage-specific regulator TpMig3, we redefine carbon catabolite repression as a central coordinating system that balances growth, energy production, and enzyme secretion. These findings provide a conceptual framework for rational engineering of filamentous fungi as industrial bio-factories and highlight the evolutionary plasticity of global regulatory networks.

## Data Availability

All data supporting the findings of this study are provided in the Supplementary Information. The raw dataset generated in the study is available on Figshare, an open data publishing platform (10.6084/m9.figshare.31341004). Any additional information required to reanalyse the data reported in this study is available from the lead contact upon request.

## Ethics Statement

All animal procedures used for antibody production were conducted in accordance with institutional guidelines for the care and use of laboratory animals

## Authors’ contribution

AR and SSY conceptualised the study and designed experiments; AR did gene deletion, molecular biology work and biochemical assays; RK cloned and purified 6X-His tagged Mig1 and Mig3 Zn fingers; AR and RK carried out antibody generation in mice at the in-house animal facility at ICGEB, New Delhi; RNA-seq data were analysed by AR and MG. AR conducted EMSA, BLI, chemical genetic experiments, ChIP-seq and proteomics sample preparation, data generation and computational analysis; AR and SSY wrote the manuscript. All authors read and approved the final manuscript.

## Supporting information

Supplemental Table - Primers used in the study

## Acknowledgements

We thank Prof Vera Meyer, Technische Universität Berlin, Germany, for providing pFW15.1 and pFW9.3 for constructing a conditional Tet-off system for *Talaromyces pinophilus*. We also thank Dr Naseem Gaur, ICGEB, New Delhi, for his guidance in carrying out EMSA. We acknowledge Girish H Rajacharya for managing the mass spectrometry facility at ICGEB, New Delhi. AR was supported by DST-SERB, India, via a national postdoctoral fellowship (PDF/2018/002549). This work was funded by the Department of Biotechnology, Government of India, via Bioenergy Centre grant no. BT/PR/Centre/03/2011-Phase-II. English language editing was assisted by an AI-based tool (ChatGPT, OpenAI); no scientific content was generated using AI.

## Copyright and Permissions Statement

The authors confirm that this manuscript contains only original material created by the authors. Therefore, no additional copyright permissions were sought.

