## Supplementary file for "Synthetic lethality of fungal catabolite repressors reveals essential coupling of carbon repression with mitochondrial and ribosomal gene expression"

Disrupting a novel C2H2 Zinc finger catabolite repressor PfMig3 is synthetic lethal with the deletion of global catabolite repressor PfMig1 in hypercellulolytic fungus Penicillium funiculosum NCIM1228

Supplementary File

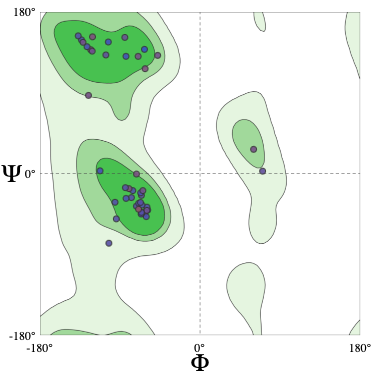

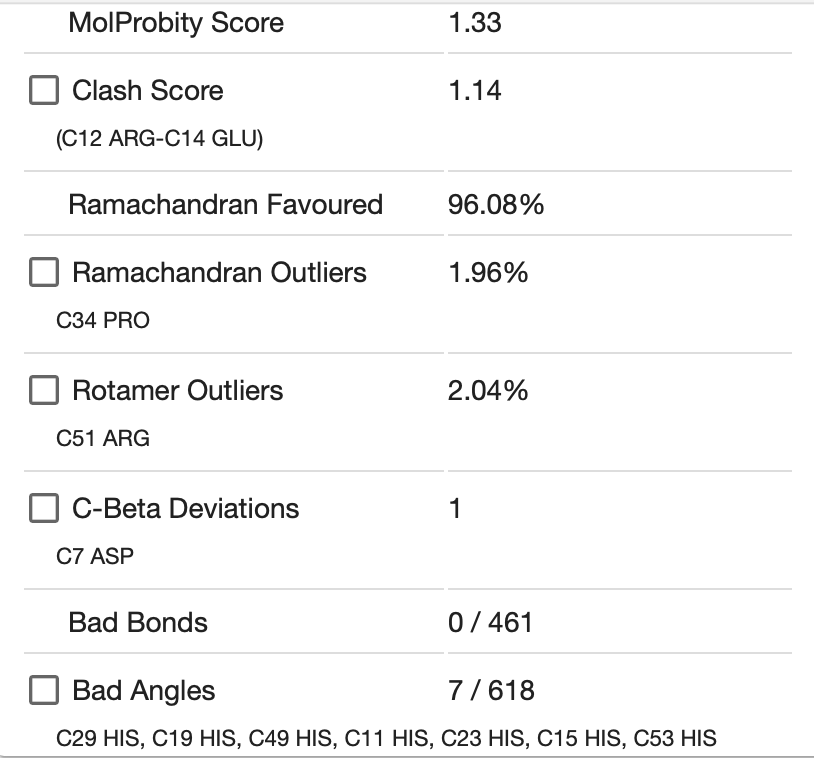

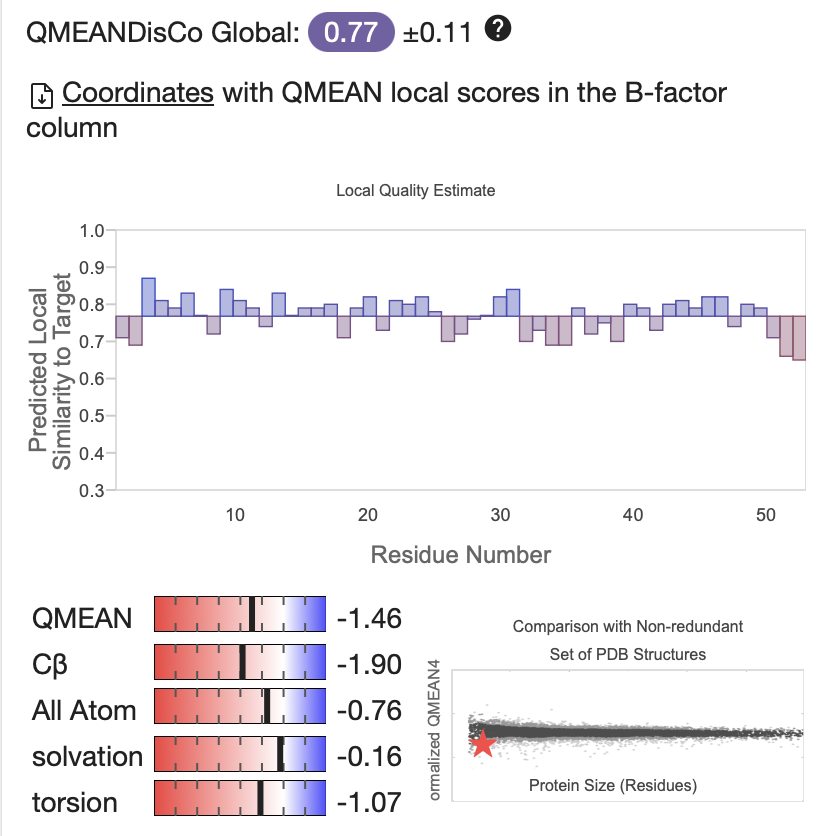

**Mig1**

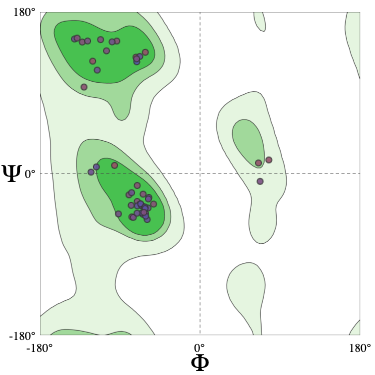

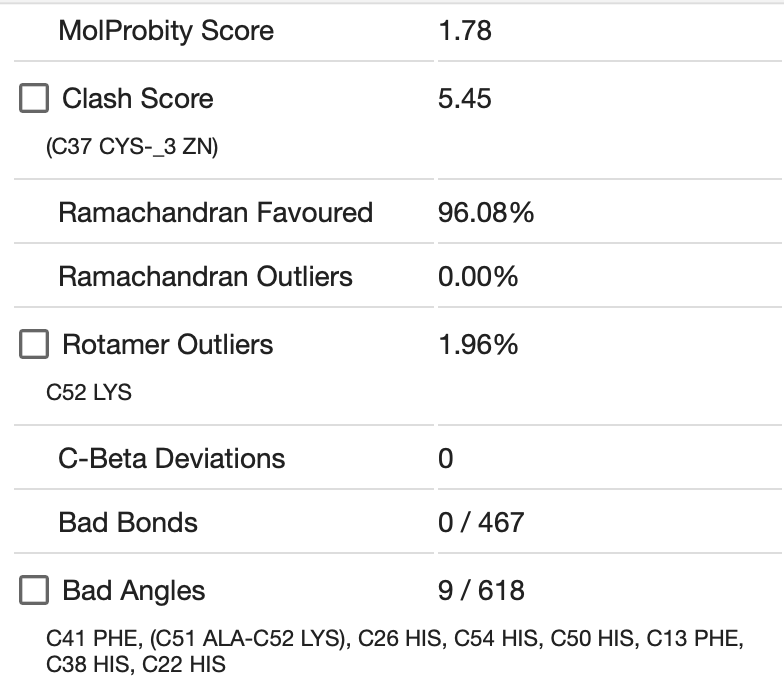

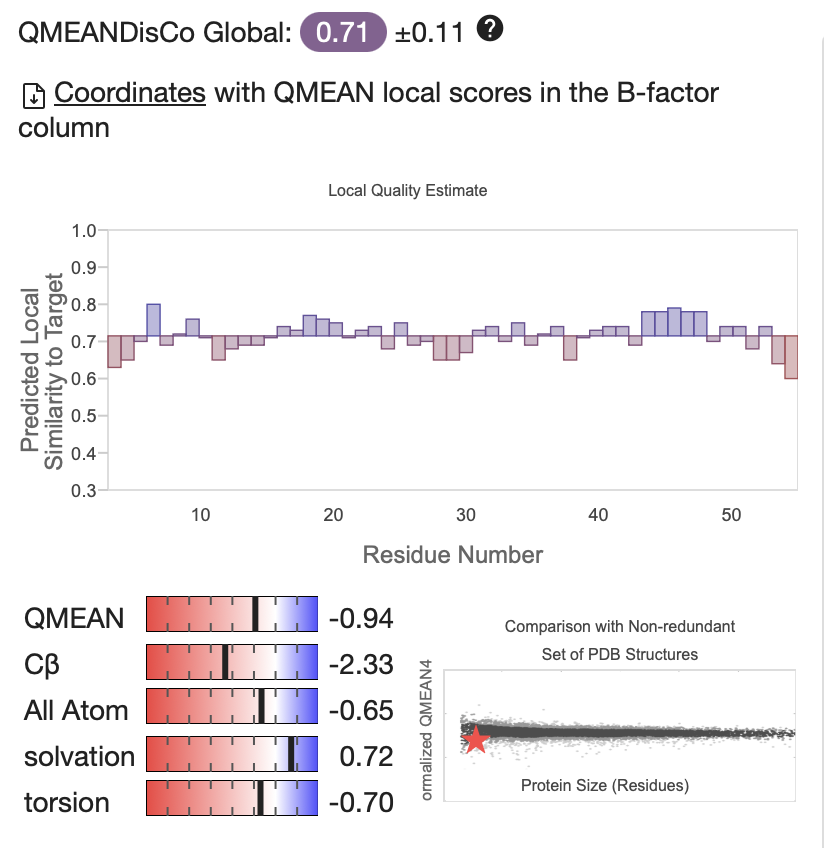

**Mig3**

Fig. S1. Ramachandran plot and coordinates of the SWISS-Model of PfMig3 and PfMig1.

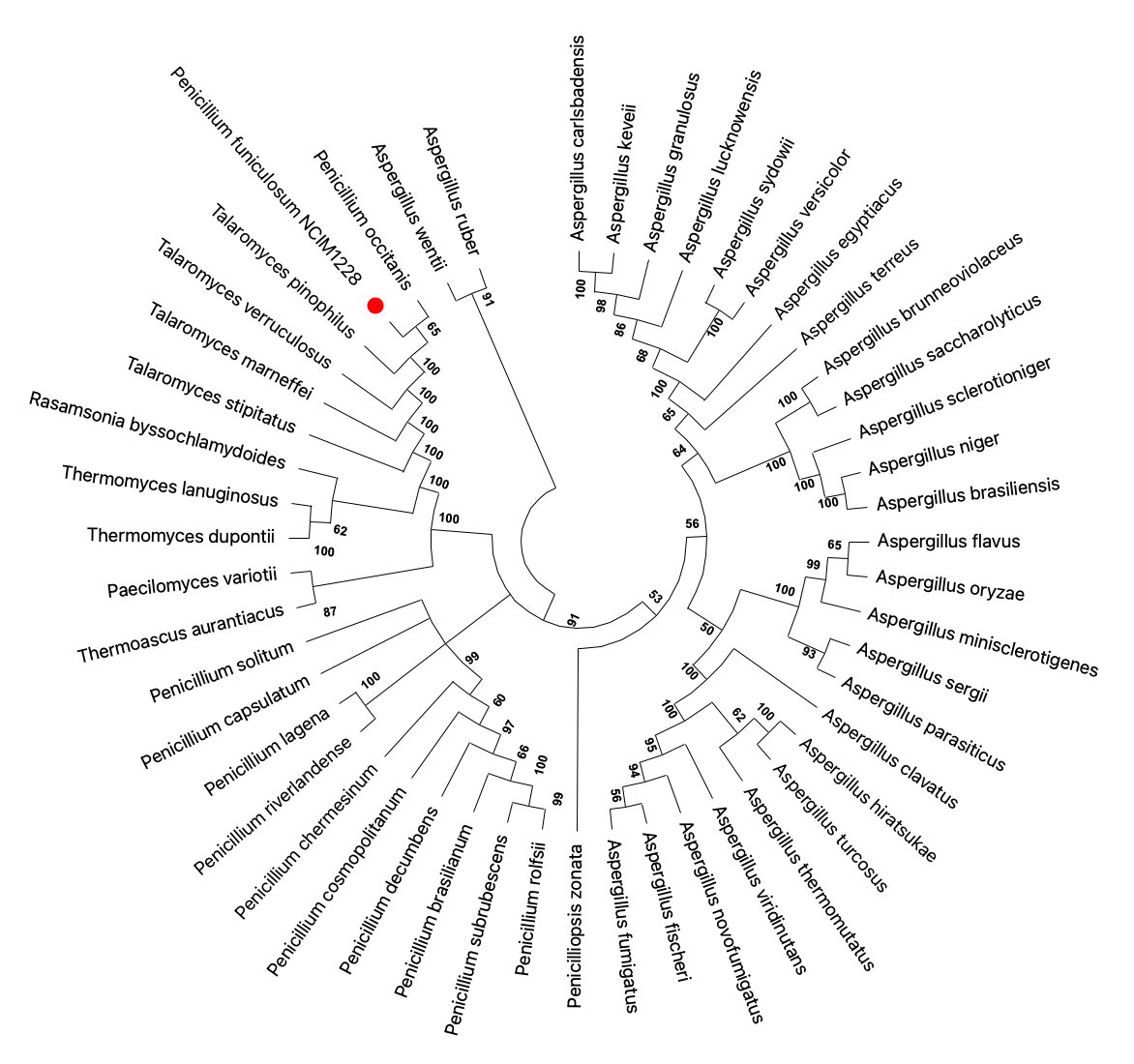

**Fig. S2. Phylogenetic tree of TpMig3 homologs.** Protein sequences of TpMig3 homologs from 50 fungal species representing the Eurotiales order were taken to construct the phylogenetic tree. TpMig3 shares one of the most recently evolved clades along with *Talaromyces pinophilus* and *Penicillium marneffei*.

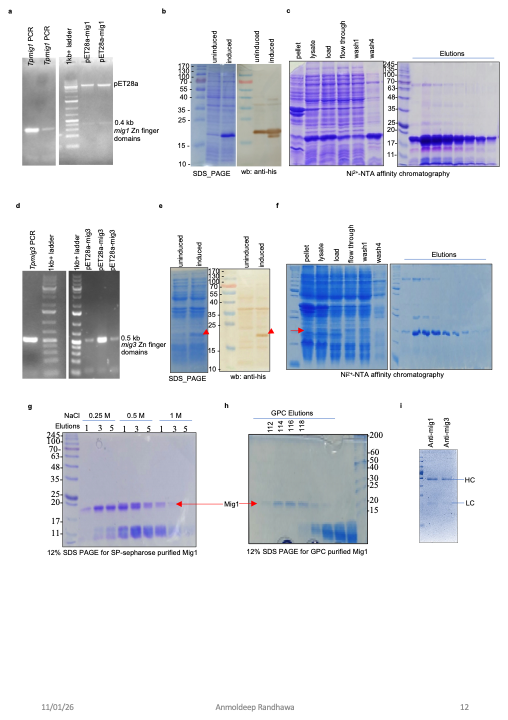

**Fig. S3. a,** PCR amplification of Mig1 gene fragment encoding for zinc fingers, and right panel shows the restriction digestion of pET28a-6xhis-Mig1. **b,** SDS-PAGE and anti-His western blotting of BL21cell lysates expressing Mig1 before and after induction. **c,** SDS-PAGE exhibiting steps of Ni-NTA affinity chromatography to achieve purified Mig1. **d,** PCR amplification of Mig3 gene fragment encoding for zinc fingers, and right panel shows the colony PCR of *E. coli* having pET28a-6xhis-Mig3. **e,** SDS-PAGE and anti-His western blotting of BL21cell lysates expressing Mig3 before and after induction. **f,** SDS-PAGE exhibiting steps of Ni-NTA affinity chromatography to achieve purified Mig3. **g,** SDS-PAGE exhibiting elution steps of SP-sepharose Cation exchange chromatography to purify Mig1. h, SDS-PAGE exhibiting elution steps of gel permeation chromatography to purify Mig1. **i,** SDS-profile of purified Anti-mig1 and Anti-mig3 antibodies; protein A/G affinity purification was conducted to purify antibodies from sera.

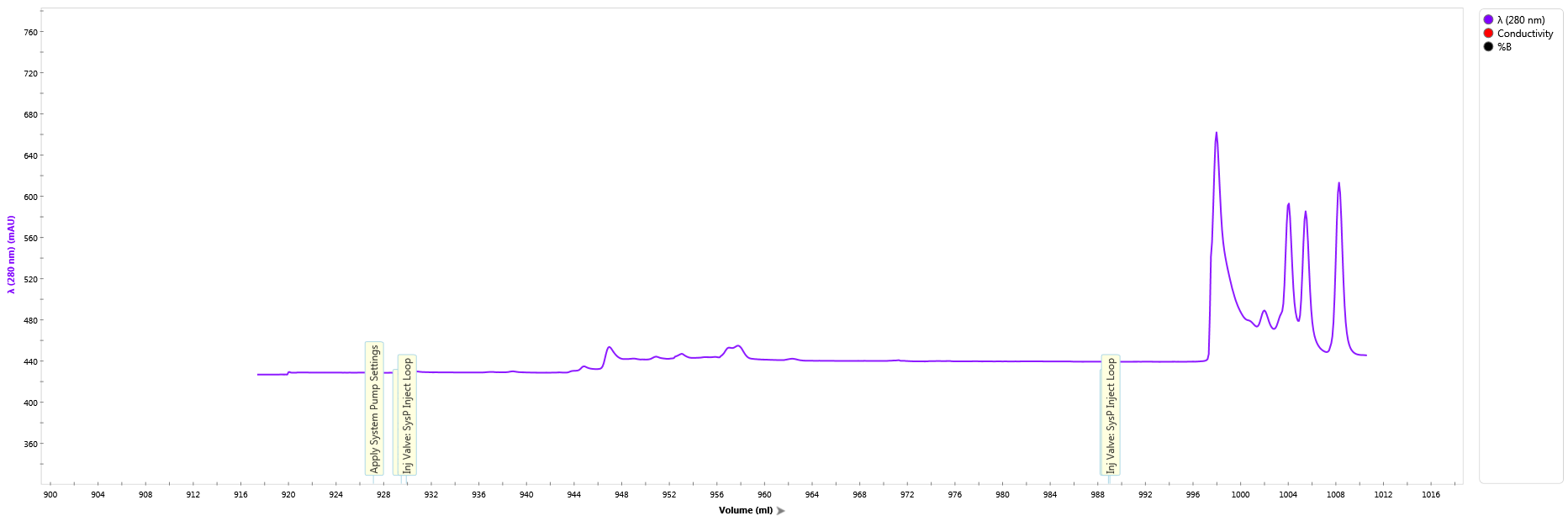

Blue dextran

RNase A (13.7 kDa)

Ovalbumin (42.7 kDa)

BSA (66.5 kDa)

**6X His Mig1 (15.5 kDa)**

**19.0**

**16.5**

**17.0**

**15.0**

**Fig. S4.** Gel permeation chromatogram of Mig1 showing dimer and polymeric forms of Mig1 in BLI buffer.
